# Regulation of the macrolide resistance ABC-F translation factor MsrD

**DOI:** 10.1101/2021.11.29.470318

**Authors:** Corentin R. Fostier, Farès Ousalem, Elodie C. Leroy, Saravuth Ngo, Heddy Soufari, C. Axel Innis, Yaser Hashem, Grégory Boёl

## Abstract

<u>A</u>ntibiotic <u>r</u>esistance ABC-Fs (ARE ABC-Fs) are translation factors currently proliferating among human pathogens that provide resistance against clinically important ribosome-targeting antibiotics. Here, we combine genetic and structural approaches to determine the regulation of *streptococcal* ARE ABC-F gene *msrD* in response to macrolide exposure and also demonstrate that MsrD twin-ATPase sites work asymmetrically to mediate the dynamic of MsrD interaction with the ribosome. We show that cladinose-containing macrolides lead to insertion of MsrDL leader peptide into an undocumented conserved crevice of the ribosomal exit tunnel concomitantly with 23S rRNA rearrangements that prevent peptide bond formation and preclude accommodation of release factors. The stalled ribosome obstructs formation of a Rho-independent terminator which prevents *msrD* transcriptional attenuation. This stalled ribosome is rescued by MsrD, but not by MsrD mutants which do not provide antibiotic resistance, showing evidence of equivalence between MsrD function in antibiotic resistance and its action on this complex.

## INTRODUCTION

ABC-F ATPases belonging to the <u>A</u>TP-<u>B</u>inding <u>C</u>assette (ABC) superfamily are translation factors and some of them, termed <u>a</u>ntibiotic <u>r</u>esistance ABC-Fs (ARE ABC-Fs), confer resistance to clinically important antibiotics that bind to the ribosomal <u>p</u>eptidyl <u>t</u>ransferase <u>c</u>enter (PTC) and/or the <u>n</u>ascent <u>p</u>eptide <u>e</u>xit <u>t</u>unnel (NPET)^1–7^. ABC-F proteins are composed of two ABC domains (or <u>N</u>ucleotide <u>B</u>inding <u>D</u>omains, NBDs) joined by a linker region called <u>P</u>-site <u>t</u>RNA-<u>i</u>nteraction <u>m</u>otif (PtIM)^8,9^, also termed <u>a</u>ntibiotic <u>r</u>esistance <u>d</u>eterminant (ARD) for ARE ABC-Fs^1–3^. The two ABC domains dimerize after binding of two ATP molecules and in this conformation the factor can bind the ribosomal E-site^9^, where the PtIM adopts an a-helical hairpin conformation that interacts directly with the peptidyl-tRNA and extends toward the PTC/NPET. Three antibiotic resistance phenotypes are associated with ARE ABC-Fs: (i) MKS_B_ (<u>M</u>acrolides, <u>K</u>etolides, <u>S</u>treptogramins group <u>B</u>); (ii) PLS_A_ (<u>P</u>leuromutilins, <u>L</u>incosamides, <u>S</u>treptogramins group <u>A</u>); (iii) PhO (<u>Ph</u>enicols, <u>O</u>xazolidinones)^10,11^. However, despite structural investigations^1–3,5,7^, the exact molecular mechanism of action of ARE ABC-Fs remains unclear.

Over the last forty years, biochemical and structural investigations demonstrated the ability of elongating <u>n</u>ascent <u>c</u>hains (NC) into the ribosomal tunnel to interact with metabolites or antibiotics, thus adapting protein synthesis to environmental cues^12–16^. In human pathogens, antibiotic-dependent formation of <u>s</u>talled <u>r</u>ibosome <u>c</u>omplexes (SRCs) on regulatory ORFs, named leader peptides, can subsequently allow regulation of downstream resistance genes in response to antibiotic exposure^17–20^.

ARE ABC-F gene *msrD* is a member of the *msr* (<u>m</u>acrolide and <u>s</u>treptogramin B <u>r</u>esistant) gene group previously referred as *mel*^21,22^. This gene is generally found in operon with a macrolide efflux facilitator *(mefA* or *mefE) gene,* the two corresponding proteins acting synergistically to confer macrolide resistance^23–26^. This operon is part of the <u>M</u>acrolide <u>E</u>fflux <u>G</u>enetic <u>A</u>ssembly (MEGA)^27,28^ that disseminates in human pathogens on mobile genetic elements: conjugative transposons like the tn916-type^29^ and conjugative prophages like the Φ1207.3^30^ among *Streptococci,* conjugative plasmids like the pMUR050 ^31^ among

*Enterobacteria.* In *Streptococci,* the operon is transcribed in presence of erythromycin (ERY) as a polycistronic mRNA from a single promoter located ~350 bp upstream *mefA^32,33^.* It is also regulated by ribosome-mediated transcription attenuation via MefAL leader peptide (MTASMRLR), which is closely related to ErmDL leader peptide (MTHSMRLRFPTLNQ), both polypeptides harboring the characteristic macrolide-arrest motif Arg/Lys-X-Arg/Lys so called “+x+”^14,33–35^.

Here, we report the presence of a second transcriptional attenuator on the *mefA/msrD* operon regulating exclusively *msrD* expression upon macrolide exposure. Presence of ERY induces ribosome stalling on the previously identified *msrDL* leader peptide (encoding MYLIFM)^34^, allowing <u>RNA</u> <u>p</u>olymerase (RNAP) to bypass an intrinsic terminator. Our findings demonstrate stalling occurs due to hampered translation termination on UAA stop codon, action of both <u>r</u>elease <u>f</u>actors <u>1</u> and <u>2</u> (RF1 and RF2) being inhibited. Our results are supported by the cryo-EM structure of MsrDL-SRC that provides molecular insight on how the PTC precludes productive accommodation of RF1/RF2 and how the NC discriminates tunnel-bound cladinose-containing macrolides. The path of the NC within the tunnel greatly differs from previously described leader peptides^12–16^ with the NC latching into a ribosomal crevice, conserved from prokaryotes to eukaryotes, delimited by 23S rRNA nucleotides U2584, U2586, G2608 and C2610 that may form a novel ribosomal functional site. Finally, our results demonstrate that the two ATPase sites of MsrD perform different functions and the protein can negatively self-regulate its synthesis in presence of ERY by direct action on MsrDL-stalled ribosomes.

## RESULTS

### MsrD provides macrolide and ketolide resistance by direct interaction with ribosome

The *mefA/msrD* macrolide resistance operon (Fig. 1a) is part of the MEGA element currently spreading among clinical isolates and livestock^33,36–38^. A phylogenetical analysis (Supplementary fig. 1a) shows that the operon disseminates predominantly in Gram-positive firmicutes (mostly *Streptococci*) and in some Gram-negative proteobacteria (such as *Haemophilus influenzae, Neisseria gonorrhoeae or Escherichia coli).* Gene *msrD* shares ~62 % identity with its closest homolog *msrE*^7^, both factors exhibiting the canonical ABC-F organization (Fig. 1b and Supplementary fig. 1b).

**Figure 1.**
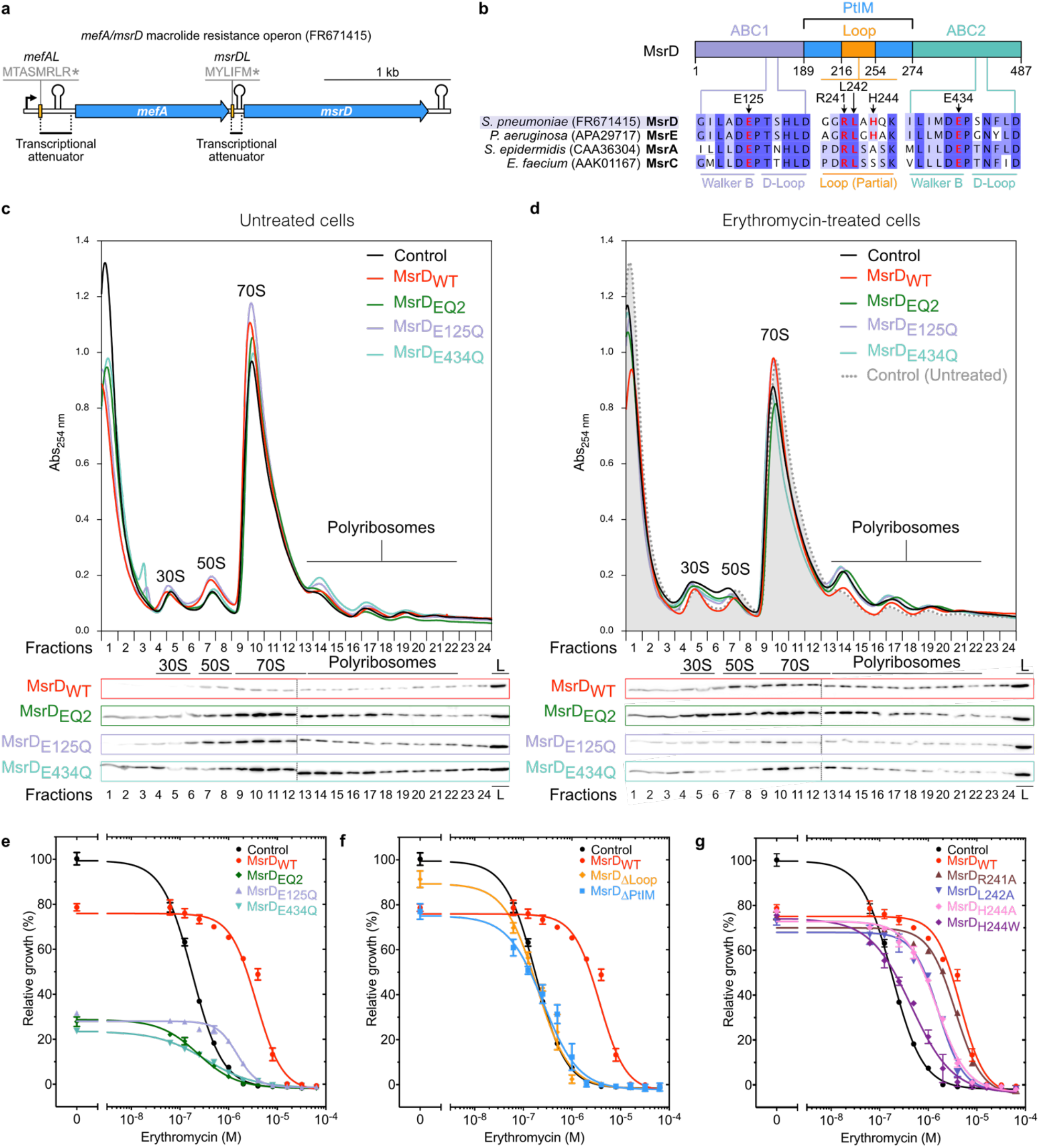
Translation factor MsrD alleviates erythromycin effects upon translation *in vivo*. (a) Organization of the *mefA/msrD* macrolide resistance operon. In presence of ERY, ribosomes stall during translation of *mefAL* leading to transcription anti-attenuation, both *mefA* and *msrD* being then transcribed. Similar mechanism occurs during translation of *msrDL,* which regulates transcription of *msrD* only. (b) Sequence alignment of various *msr* homologs visualized with Jalview according to percentage of identity. Positions of tested mutations are highlighted in red and indicated by arrows. Main features of ABC-F proteins are indicated on the schematic. Genebank accession numbers are indicated between brackets. See also Supplementary Fig. 1b. (c and d) Polyribosomes analysis and western blotting of *E. coli* DB10 expressing a control (pBAD-*Control*), *msrD_WT_* (pBAD-*msrD*_WT_), *msrD*_EQ2_ (pBAD-*msrD*_EQ2_), *msrD*_E125Q_ (pBAD-*msrD_E125Q_*) or *msrD_E434Q_* (pBAD-*msrD_E434Q_*) untreated or treated for 1 h with 25 μM ERY during mid-log phase. “L” stands for total lysate. See also Supplementary fig. 1c and 1d and Methods. (e to g) Relative growth of *E. coli* DB10 expressing *msrD* variants in presence of ERY after 24 h. Optical densities were normalized relative to optical densities of *E. coli* DB10 pBAD-*Control* grown in the absence of ERY. Error bars represent mean ± s.d. for triplicate experiments. See also supplementary Table 1.

Model bacteria *Escherichia coli (E. coli)* presents undeniable advantages for genetic and molecular biology, but presence of multidrug efflux systems greatly limits its use in antibiotic research^39,40^. To circumvent this limitation, we took advantage of *E. coli* DB10 strain which exhibits exacerbated sensitivity toward macrolide antibiotics^41^. Heterologous expression of *msrDWT* from an arabinose-inducible promoter conferred macrolide and ketolide resistance phenotype to *E. coli* DB10 strain (Table 1), demonstrating the functionality of the factor in this organism. Expression of *msrDWT* resulted in 8-fold <u>m</u>inimal <u>i</u>nhibitory <u>c</u>oncentration (MIC) increase for 14-membered macrolide ERY, 8-fold for 15-membered macrolide azithromycin (AZI) and 2-fold for telithromycin (TEL, ketolide antibiotic derived from 14-membered macrolides). Similar results were reported for *Streptococcus pneumoniae* clinical isolates^23^. However, no change in MIC was observed for 16-membered macrolides tylosin (TYL) and spiramycin (SPI), neither for non-macrolide antibiotics lincomycin (LNC), linezolid (LNZ) and retapamulin (RTP) (Table 1).

**Table 1.** Minimum inhibitory concentration (MIC) and half maximal inhibitory concentration (IC_50_) of *E. coli* DB10 expressing *msrD* in presence of ribosome-targeting antibiotics. See Methods for experimental details.

|  | **pBAD-Control** |  | **pBAD-*msrD*<sub>WT</sub>** |  |
| --- | --- | --- | --- | --- |
| **Antibiotic** | **MIC (μM)** | **IC<sub>50</sub> (μM)** | **MIC (μM)** | **IC<sub>50</sub> (μM)** |
| **Erythromycin** | 2 | 0,179 ± 0,007 | 16 | 4,592 ± 0,582 |
| **Azithromycin** | 0.25 | 0,046 ± 0,002 | 2 | 0,953 ± 0,053 |
| **Telithromycin** | 1 | 0,086 ± 0,008 | 2 | 0,714 ± 0,066 |
| **Tylosin** | 4 | 1,649 ± 0,151 | 4 | 1,672 ± 0,103 |
| **Spiramycin** | 2 | 0,758 ± 0,063 | 2 | 0,81 ± 0,078 |
| **Lincomycin** | 32 | 12,61 ± 2,154 | 32 | 12,62 ± 1,27 |
| **Linezolid** | 32 | 13,62 ± 2,064 | 32 | 13,41 ± 1,137 |
| **Retapamulin** | 1 | 0,071 ± 0,004 | 1 | 0,075 ± 0,007 |

Comparison of polysome profiles from bacteria unexposed or exposed to ERY show a stabilization of ribosome disomes (Fig. 1c, 1d and Supplementary fig. 1c) as expected for an antibiotic that inhibits ribosome elongation. The disome accumulation likely results from elongating ribosomes that collide on ERY-stalled ribosomes arrested at the beginning of the coding sequence. In presence of ERY ribosomes preferentially stall when they encounter “+x+” motifs which are present in the first 30 amino-acids of ~25% of *E. coli* proteins^42,43^. Expression of *msrD*_WT_ alleviates the accumulation of disomes and restores a profile identical to that without ERY (Fig. 1d versus 1c and Supplementary fig. 1c). Western blots of the polysome fractions show an MsrDWT association with the 70S ribosome and the 50S subunit and polysomes (Fig. 1d and Supplementary fig. 1c). These results agree with a model in which MsrD_WT_ directly rescues ERY-stalled ribosomes and that MsrD can restore normal translation in presence of the antibiotic.

To understand the function of the ATPase activity of MsrD, we constructed ATPase-deficient variants MsrD_E125Q_, MsrD_E434Q_ and MsrD_E125Q/E434Q_ (hereafter referred as MsrD_EQ2_) by replacing the catalytic glutamic acid residue (E) in Walker B motif of each or both ABC domain by a glutamine (Q) (Fig. 1b and Supplementary fig. 1b). This mutation strongly reduces the ATP hydrolysis while preserving the local stereochemistry of active site and has been extensively used to trap ABC-F factors on the ribosome in ATP-bound conformation^1–3,5,9^. Expression of *msrD*_EQ2_ in absence of ERY inhibits cell growth after OD_600_=0.4 and shows an accumulation of disomes on polysome profile (Supplementary fig. 1c) and western-blotting reveals its association with the 30S subunit, 70S ribosome and polyribosomes (Fig. 1d and Supplementary fig. 1c). These results shows that MsrD_EQ2_ binds to the ribosome and inhibits its elongation as observed for most of the ABC-F carrying the E-to-Q mutation^8,44^. In presence of ERY, the disome and trisome accumulation increases and western blots show an increase of MsrD_EQ2_ in the disome fraction (Fig. 1d and Supplementary fig. 1c). Still, the ATPase activity of MsrD impacts its binding preferences, MsrD_WT_ interacting more with the 50S subunit in contrast to MsrD_EQ2_ interacting more with the 70S ribosome (Fig. 1d and Supplementary fig. 1c). Interestingly, the MsrD_E125Q_ variant has an interaction profile with the ribosome similar to MsrD_WT_ whether MsrD_E434Q_ has one closer to MsrDEQ2.

A model consistent with these observations would imply that MsrD_WT_ binds to ERY-stalled ribosomes in an ATP-bound close conformation without the need of its ATPase activity, then ATPase activity triggers 70S dissociation, MsrD_WT_ staying associated with the 50S subunit. Similar results were observed by expressing *msrD*_WT_ and *msrD*_EQ2_ in *E. coli* K12 MG1655 (Supplementary fig. 1d) in absence of ERY and with chloramphenicol being added to stabilize polysomes. They also show a decrease of the 70S peak correlated with an increase of 30S and 50S peaks for *msrD*_WT_ condition when compared to the control, supporting the proposed dissociation model. western blots show more MsrD_EQ2_ than MsrD_WT_ in soluble lysates (Supplementary fig. 1c) but MsrD_EQ2_ inhibits protein synthesis including its own synthesis. Therefore, we hypothesized that MsrD_EQ2_ is less expressed but more soluble, which has been confirmed with a solubility test (Supplementary fig. 1e).

We evaluated the influence of E-to-Q mutations of MsrD on bacterial growth and antibiotic resistance phenotype. All the mutations impair bacterial growth and antibiotic resistance phenotype (Fig. 1e), while MsrD_E125Q_ and MsrD_EQ2_ have a similar effect on bacterial growth in absence or at low ERY concentration, MsrD_E434Q_ is more toxic. However, MsrD_E125Q_ variant maintains some resistance (MIC = 4 μM) compared to control (MIC = 2 μM) which is also supported by the fact that MsrD_E125Q_ IC_50_ is ~8 times higher than Control IC_50_ (Fig. 1e and Supplementary Table 1). These results together with polysome profile experiments (Fig. 1c, 1d and Supplementary fig. 1d) suggest that interaction of MsrD with the 50S is necessary for antibiotic resistance mechanism.

The two ATP hydrolysis sites (Supplementary fig. 1b) have therefore different tasks and are functionally asymmetric, ATP site II achieving in part the mechanism of ERY-stalled ribosome rescue. We also evaluated the importance of other residues of MsrD. Truncation of the PtIM (replaced by three glycine) or just the Loop (ΔPtIM, ΔLoop respectively) completely abolished the resistance phenotype *in vivo* (Fig. 1f) as reported *in vitro* for MsrE^7^. Punctual mutations in the Loop (Fig. 1b) affect the resistance phenotype to a different extent, mutation of R241 into alanine has almost no effect on the phenotype, but equivalent mutation for L242 or H244 reduces IC_50_ by ~3 fold (Fig. 1g and Supplementary Table 1), demonstrating that these residues are important for MsrD function, but not essential even if they are predicted at the vicinity of the antibiotic (Supplementary fig. 1f) in the MsrE-ribosome complex^7^. Interestingly, this finding contrasts with mutagenesis assays for MsrE showing that mutation R241A reduces the resistance by more than 50%. Replacement of H244 by the larger residue tryptophan should displace the drug by steric occlusion, but the variant lost most of its antibiotic resistance phenotype (Fig. 1g). Similarly, MsrD_WT_ does not provide resistance to TYL nor SPI (Table 1), while both antibiotics should clash with MsrD according to MsrE structure (Supplementary fig. 1f). Same observation can be made with LNC, LNZ or RTP (Table 1 and Supplementary fig. 1f). Therefore, MsrD action on the antibiotic is indirect and direct steric occlusion to displace the antibiotic seems excluded.

### Ribosome-mediated transcriptional attenuation regulates *msrD* expression

Analysis of the *mefA/msrD* operon sequence exposes a stringent <u>r</u>ibosome <u>b</u>inding <u>s</u>ite (RBS, whose sequence is 5’-GGAGGA-3’) only 8 bp downstream *mefA* stop codon and possibly required for translation of the downstream putative small ORF *msrDL* (Supplementary fig. 2a). In order to test the influence of *msrDL* on *msrD* expression, a fluorescent reporter, which contains the sequence spanning from the first nucleotide downstream *mefA* stop codon to *msrD* three first codons fused to the <u>Y</u>ellow <u>F</u>luorescent <u>P</u>rotein gene (YFP), was cloned in the low copy pMMB plasmid^45^ under the control of IPTG-inducible P_LlacO-1_ promoter^46^. The resulting plasmid, *pMMB-msrDL-msrD_(1-3)_:yfp,* was used to directly follow the fluorescence reflecting *yfp* expression (Fig. 2a).

**Figure 2.**
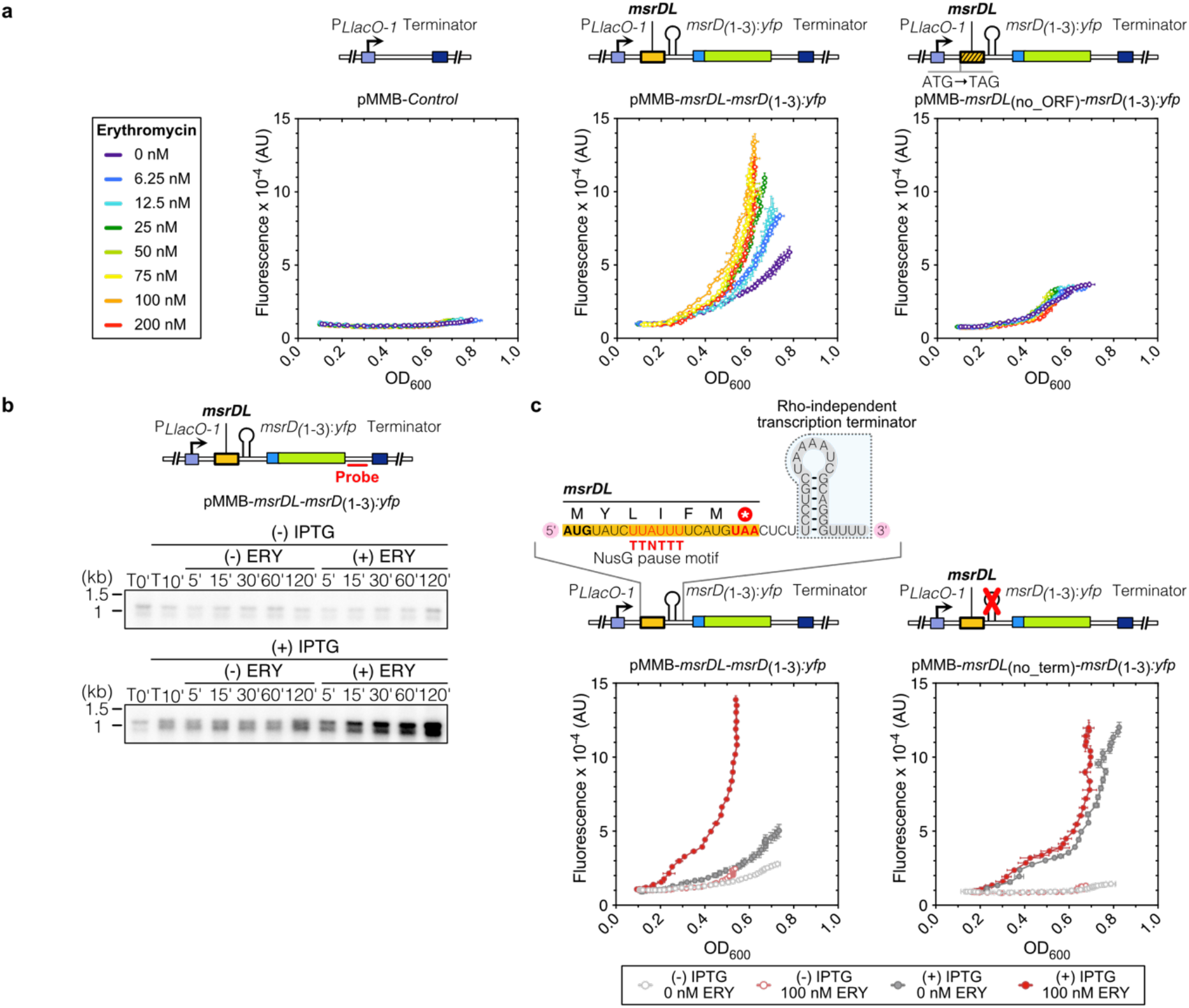
Erythromycin-dependent transcriptional attenuation regulates *msrD* expression. (a) ERY-dependent induction of *msrD*_(1-3)_:*yfp*. Fluorescent reporters shown on schematics have been introduced in *E. coli* DB10 and grew in presence of 1 mM IPTG and increasing sublethal ERY concentrations during 17 h. Fluorescence has been plotted against OD_600_, error bars for both axes represent mean ± s.d. for triplicate experiments. (b) Northern analysis of *msrD* transcript. RNAs were extracted before adding or not 1 mM IPTG (T_0_’), 10 min after adding IPTG (T_10_’), and 5, 15, 30, 60, 120 min after adding or not 100 nM ERY. Location of probe in *msrD*_(1-3)_:*yfp* 3’ UTR is shown on the schematic. The presence of a second band was also observed but we hypothesized that it was abortive transcript resulting from the construct insofar as its presence correlates with induction by IPTG and ERY, and is no longer detectable using other probes (data not shown). Note the presence of leaky transcription in absence of IPTG, that is slightly amplified in presence of ERY. (c) Deletion of the intrinsic terminator between *msrDL* and *msrD*_(1-3)_:*yfp* leads to a constitutive induction in absence of ERY. Error bars for both axes represent mean ± s.d. for triplicate experiments.

Basal fluorescence level was found stable in *E. coli* DB10 strain containing pMMB-*Control* independently of ERY concentration (Fig. 2a) while a fluorescence increase correlated with ERY concentration was observed in cells containing pMMB-*msrDL-msrD*_(1-3)_:*yfp*, indicating an ERY-dependent induction. Optimal induction was found at 100 nM ERY while only 6.25 nM was sufficient to significantly induce *msrD*_(1-3)_:*yfp* expression, demonstrating the high sensitivity of the system toward its inducer. A reduced, but significative fluorescence signal was detected at 0 nM ERY indicating an imperfect repression of the regulation. Inactivation of *msrDL* ORF by replacement of the start codon by an amber stop codon (pMMB-*msrDL*_(no_ORF)_*-msrD*_(1-3)_:*yfp*) resulted in the elimination of the fluorescence induction (Fig. 2a), demonstrating that *msrDL* translation is an important and necessary cis-acting feature to regulate *msrD_(1-3)_:yfp* expression as reported for *ermC*^47^.

In bacteria, antibiotic-dependent gene induction relies mainly on translational or transcriptional attenuation^48^. To determine if *msrDL* prompts a transcriptional or translational attenuation, northern blots analysis with a probe located in *msrD*_(1-3)_:*yfp* 3’ UTR were performed on total RNA extracted from cell exposed or not to ERY and/or IPTG. A 1.2-kb band corresponding to *msrDL*-*msrD*_(1-3)_:*yfp* mRNA is transcribed in presence of IPTG and starts to increase steadily 5 min after exposure to 100 nM ERY (Fig. 2b). Consistent with this ERY-dependent transcript accumulation, we concluded that *msrD* regulation by *msrDL* occurs at the transcription level.

Transcription attenuation employs premature transcription termination via: (i) <u>R</u>ho-<u>d</u>ependent <u>t</u>erminator (RDT) by binding of Rho factor, a RecA-type ATPase, to a C-rich and G-poor sequence known as Rho utilization site^49^; (ii) <u>R</u>ho-<u>i</u>ndependent <u>t</u>erminator (RIT) where a stable GC-rich stem-loop followed by a poly-uridine sequence causes RNAP to drop off^50^. Treatment with bicyclomycin, a Rho inhibitor, did not result in constitutive expression of the reporter, thus excluding the possibility of RDT attenuation (Supplementary fig. 2b). Analysis of the intergenic region between *msrDL* and *msrD* using ARNold server^51^ revealed the presence of a RIT with a ΔG of −7.9 kcal.mol^-1^. We also identified a consensus TTNTTT NusG-dependent RNAP pausing site embedded in *msrDL* sequence (Fig. 2c and Supplementary fig. 2a), as previously described for *vmlR* ^52,53^. In order to test this possible transcription termination activity, we generated a mutant by removing the putative terminator (referred as pMMB-msrDL_(no_term)_-msrD_(1-3)_:*yfp*). Deletion of the terminator resulted in a constitutive expression of the fluorescent reporter in absence of ERY, indicating that the terminator is a necessary cis-acting regulatory feature that can terminate transcription (Fig. 2c). These results validate the presence of a second transcriptional attenuator in the *mefA/msrD* operon that regulates exclusively *msrD*.

### Selective drug-sensing by MsrDL prevents its translation elongation and termination

To determine how this regulation occurs via ribosome-mediated mechanism, translation of *msrDL* mRNA with the PURE system^54^ was subjected to various antibiotics and its propensity to form SRC was analyzed by toeprinting^55^ (Fig. 3 and Supplementary fig. 3). A strong toeprint signal was observed in presence of ERY and AZI, while it was absent in presence of TEL, TYL and SPI (Fig. 3a). This *in vitro* result is correlated to *in vivo* observations made with the *msrD_(1-3)_:yfp* reporter gene which was also only induced by ERY and AZI, but not by TEL, TYL, SPI and non-macrolide antibiotics (Fig. 3b).

**Figure 3.**
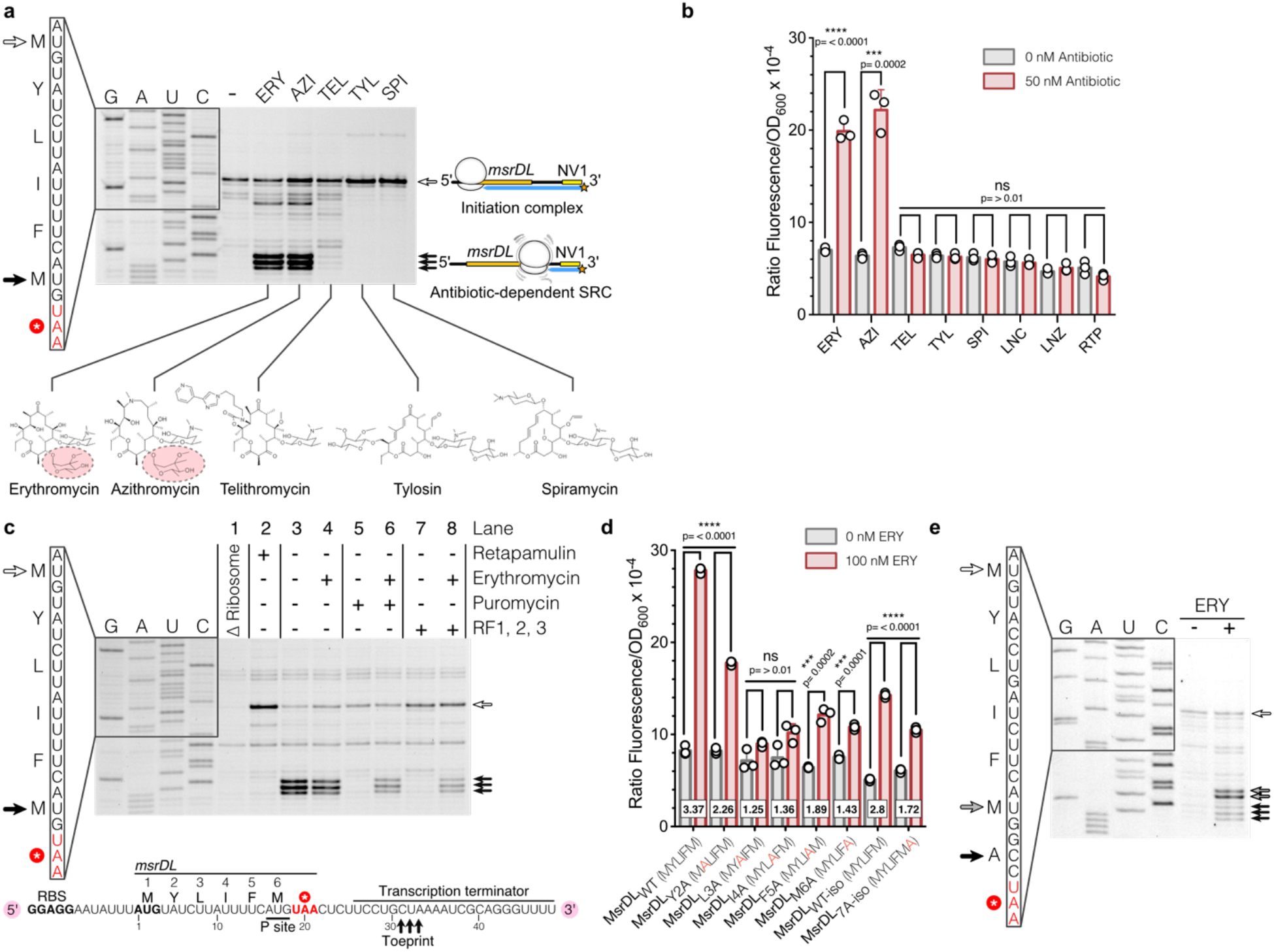
MsrDL is a macrolide-sensing nascent chain that stalls the ribosome. (a) Toeprinting assay of *msrDL* in the absence (-) or in the presence of 50 μM of various macrolide antibiotics. White arrow indicates initiation codon. Black arrows indicate ribosome stalling, with M6 codon in the P site. Chemical structure of antibiotics is shown, C3 cladinose sugar of ERY and AZI being highlighted. See also Supplementary fig. 3. (b) *In vivo* induction of *msrD_(1:3)_:yfp* by various PTC/NPET targeting antibiotics. Bacteria containing pMMB-*msrDL*-*msrD*_(1:3):_*yfp* were grown during 17 h in presence of 1 mM IPTG, in the absence (grey histograms) or in the presence of 50 nM antibiotics (red histograms). Error bars represent mean ± s.d. for triplicate experiments. (c) Toeprinting assay depleted of release factors was performed in the absence of ribosome (line 1), in presence of 50 μM RTP to assess start codon (line 2), without or with 50 μM ERY (line 3 and 4), without or with 50 μM ERY then supplemented with 100 μM puromycin (line 5 and 6), without or with 50 μM ERY in presence of RF1/RF2/RF3 (line 7 and 8). The schematic indicates position of toeprint signal on the synthetic mRNA, P site codon of MsrDL-SRC is underlined. See also Supplementary fig. 3. (d) Effects of *msrDL* variants on the expression of *msrD_(1:3)_:yfp*. Bacteria were grown during 17 h in presence of 1 mM IPTG, in the absence (grey histograms) or in the presence of 100 nM ERY (red histograms). Error bars represent mean ± s.d. for triplicate experiments, square boxes show fold of induction. (e) Toeprinting assay performed on the MsrDL_7A-iso_ construct in absence or presence of ERY. Addition of a sense codon after M6 codon leads to translational arrest with M6 codon in the P site (grey arrows) and A7 codon in the A site in presence of ERY. A faint toeprint is also observed with A7 codon in the P site (black arrows) and the stop codon in the A site.

We further characterized the translation of *msrDL* mRNA and showed that in presence of 50 μM RTP, an inhibitor of the first peptide bond formation which allows to identify ORFs start codon^56^, a clear toeprint occurred (Fig. 3c). This toeprint corresponds to a ribosome stalled with the initiating codon in the ribosomal P site and confirms that ribosomes can initiate *msrDL* translation. The toeprint observed at positions +31, +32 and +33 relative to *msrDL* 5’ end (Fig. 3a and 3c) in presence of ERY and AZI indicates an SRC located at the termination step of MsrDL synthesis with the C-terminal methionine (M6) codon in the P site and UAA stop codon in the A site. In accordance with this, reactions depleted of release factors (RF1, RF2, RF3) (Fig. 3c) show the same toeprint result. Moreover, addition of puromycin led to the loss of this toeprint signal, while preserving it in presence of ERY (Fig. 3c). Puromycin is an aminonucleoside antibiotic mimicking A site substrate tyrosyl-tRNA and causing premature chain termination. While actively translating ribosomes are sensitive to this antibiotic, stalled ones tend to be refractory^15,16,20,57,58^. The resistance of MsrDL-SRC to puromycin is consistent with an ERY-sensing NC that leads to PTC silencing and stalling. The stalled ribosome on ORF *msrDL* covers the first half of the hairpin of the RIT and therefore prevents its formation and transcription termination (Fig. 2c and Supplementary fig. 3).

The main dissimilarity between tested macrolides resides in the presence or absence of a cladinose moiety on C3 of the macrocyclic lactone ring which appears to be critical for MsrDL-SRC formation (Fig. 3a) similarly to ErmCL^20,59^. Absence of significant toeprint in presence of TEL (whose C3 cladinose sugar is replaced by a keto group) suggests that *msrDL* is translated efficiently without stalling. TYL and SPI differ from the other macrolides by the presence of disaccharide mycaminose-mycarose in place of the C5 monosaccharide desosamine. When bound to the ribosome, this extended sugar moiety protrudes into the PTC of the ribosome^60^. Absence of intermediate toeprint and increase of the toeprint corresponding to initiation complex in presence of these two drugs imply that these antibiotics stabilize the initiation complex. Previous results^60–62^ have shown that they inhibit the formation of the peptide bond, our result suggests that it occurs mainly for the first peptide bond formation (Fig. 3a).

We submitted the sequence of MsrDL to an alanine-scanning to investigate the importance of each residue and their ability to induce *msrD*_(1-3)_:*yfp* reporter, which indirectly reports the formation of an ERY-dependent SRC. The scan shows a strong reduction of inducibility for all the mutated residues with the exception Y2 (Fig. 3d). Since MsrDL is a hexapeptide, it is evident that most of the amino acids would have their importance to maintain a conformation able to sense ERY. A complete loss of inducibility has been observed for residues L3 and I4 suggesting that these two residues were directly involved in drug-sensing, which was structurally confirmed and detailed in the next section. Moreover, considering that MsrDL inhibits the action of both RF1/RF2 in presence of ERY, we mutated UAA stop codon into RF1-specific UAG and RF2-specific UGA stop codons to investigate a putative preferential inhibition as previously shown for TnaC^63^. Mutation of UAA into UAG or UGA reduces the level of inducibility of the reporter gene (Supplementary fig. 2c) demonstrating the importance of the stop codon. This observation is corroborated by our phylogenetic analysis which revealed a large prevalence of UAA stop codon among the different *msrDL* variants (Supplementary fig. 2a). Most likely the UAA stop codon has been selected by evolution to maximize *msrD* expression in presence of an inducer. Mutagenesis of *msrDL* sequence by synonymous codons (iso-codons) as described in Methods leads to a reduction of the *yfp* expression for MsrDLWT-_iso_ construct, possibly due to the lack of NusG pausing site, but the regulation is mostly preserved (Fig. 3d). Therefore, the amino acid composition of the peptide is responsible for the ribosome stalling. Addition of an extra alanine before the stop codon in MsrDL_7A-iso_ construct reduces the regulatory potential of the leader peptide (Fig. 3d). Toeprinting assay reveals that this construct has two stalling positions: one occurs with the additional A7 codon in the ribosomal A site and the second with the stop codon in the A site (Fig. 3e). Thus, MsrDL peptide seems to induce efficient ribosomal stalling by preventing elongation and termination. When both are uncoupled like in the MsrDL_7A-iso_ construct the regulation is less efficient (Fig. 3d).

### Molecular mechanism of ribosome stalling by MsrDL

To understand how MsrDL inhibits translation termination in presence of ERY, a synthetic mRNA containing a single *msrDL* copy was translated *in vitro* with purified *E. coli* DB10 ribosomes in presence of ERY as described in Supplementary fig. 4a and the resulting sample was subjected to cryo-EM. Single-particle reconstruction of MsrDL-SRC was done by refining two subclasses, one containing only P-tRNAS and one containing P- and E-tRNAs (respectively 11% and 28.1% of total particles), as described in Methods and on Supplementary fig. 4b. It resulted an initial map of 70S ribosome (EMD-13805) with an average resolution of 3.0 Å that was subjected to multibody refinement, generating reconstructions for the body and the head of the small subunit (EMD-13807 and EMD-13808) and the large subunit (EMD-13806), presenting average resolutions of 3.08, 3.3 and 2.97 Å respectively (Supplementary fig. 4b and 4c and Supplementary Table 2). The resolutions of the derived reconstructions were consistent with clear and unambiguous assignment of ribosomal proteins side chains, rRNA nucleotides and some post-transcriptional modifications (Supplementary fig. 4d to 4f). Local resolution of the codon-anticodon allowed unambiguous identification of the P-tRNA as an elongator _Met_tRNA^Met^ (Supplementary fig. 4e), which is also confirmed by the presence of distinctive elements such as N^4^-acetylcytidine at position 34 (ac^4^C34) and N^6^-threonylcarbamoyladenosine at position 37 (t^6^A37) (Supplementary fig. 4e and 4f)^64,65^, consistently with the toeprint results which assigned the M6 codon in the P site and UAA stop codon in the A site (Fig. 3c).

A clear density corresponding to ERY bound in its canonical position was found at the entrance of the NPET (Fig. 4a, 4b and Supplementary fig. 4d), allowing clear attribution of macrocyclic lactone ring, cladinose and desosamine sugars. A continuous density at the 3’ end of P-tRNA extending within the entrance of ribosome tunnel was identified as MsrDL-NC and local resolution of ~3 Å allowed to model the peptide *de novo* (Fig. 4b and Supplementary fig. 4d). It adopts a hook-like shape with its N-terminal extremity protruding into a cavity at the entrance of the tunnel delimited by 23S rRNA nucleotides U2584, U2586, G2608 and C2610 (Fig. 4b). One noticeable feature of MsrDL interaction with the ribosome is its residue Y2 that bulges out the base U2584 to form a π-stacking with the base G2583 (Fig. 4c and 4d) in place of U2584. This unique interaction should stabilize the peptide conformation, while it appears structurally significant, it is not strictly essential since replacement of the Y2 by an alanine reduces by less than 50% the induction of the reporter gene (Fig. 3d). In addition, this residue is substituted in some MsrDL variants (Supplementary fig. 2a). MsrDL does not show numerous significant electrostatic interactions with the ribosome since it is mostly composed of hydrophobic amino acids. It is likely stabilized by hydrophobic interactions.

**Figure 4.**
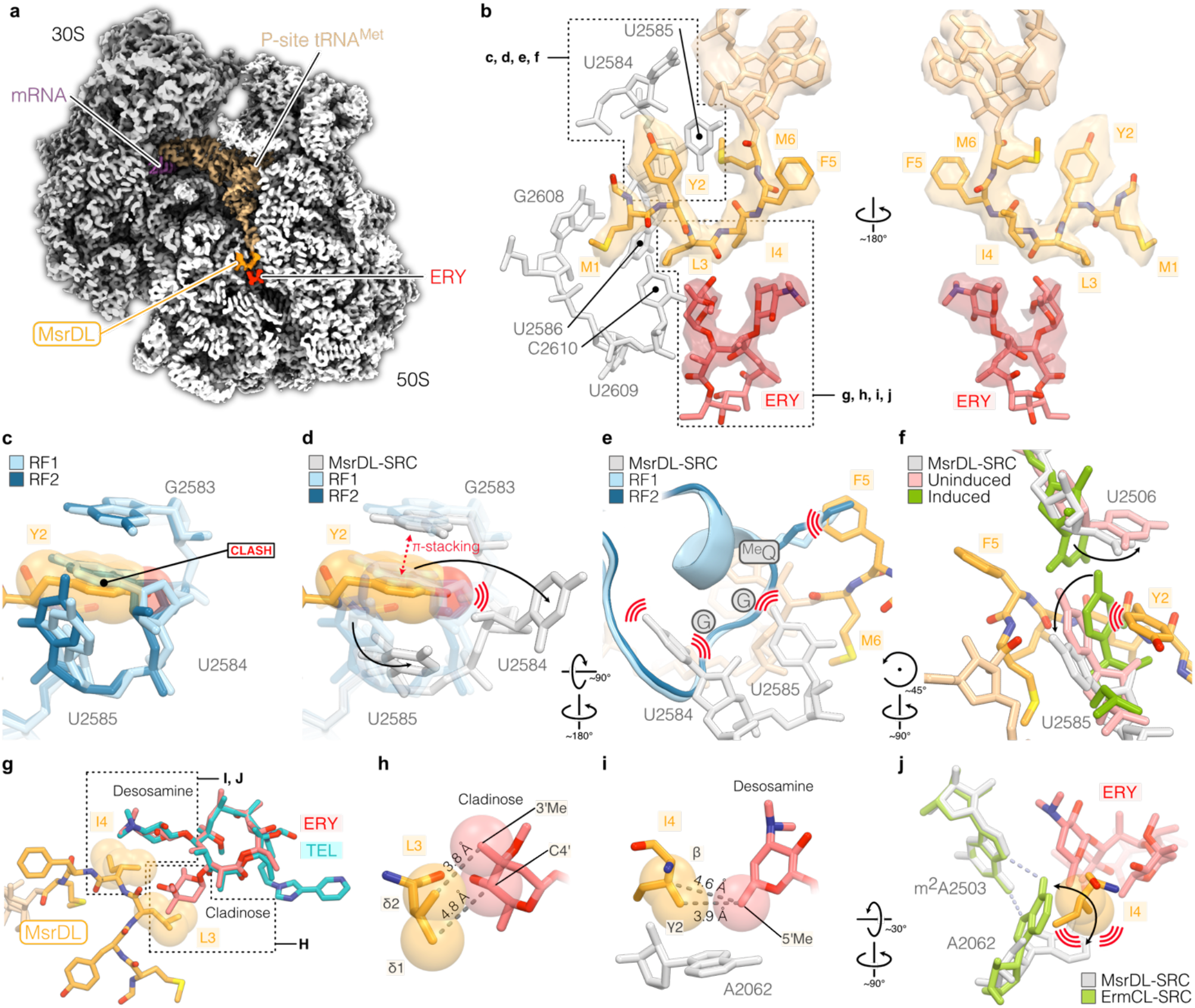
Structure of MsrDL-SRC. (a) Transverse section of the cryo-EM map showing the 30S (grey) and 50S (white) ribosomal subunits, mRNA (purple), ERY (red) and MsrDL-NC (gold) bound to P-site tRNA^Met^ (beige). (b) Close-up of MsrDL-NC within ribosomal tunnel showing experimental density and modelled structure, colored as Fig. 4A while 23S rRNA nucleotides are shown in light grey. (c and d) Presence of MsrDL residue Y2 displace nucleotide U2584 out of its position when compared to RF1-(PDB 5J30, light blue) and RF2-containing (PDB 5CZP, blue) termination complex, thus forming a π-stacking with G2583 ^66^. (e) Conformation of U2584 and U2585 prevent productive accommodation of RF1 (PDB 5J30, light blue) and RF2 (PDB 5CZP, blue) while catalytic methylated glutamine would clash with MsrDL residue F5^66^. (f) PTC in MsrDL-SRC (light grey) is stabilized in an uninduced state (PDB 1VQ6, pink) rather than in an induced state (PDB 1VQN, green) as U2585 is pushed back by MsrDL residue Y2^67^. (g to i) Molecular basis for C3 cladinose sugar recognition by MsrDL. Residue L3 is at proximity of cladinose sugar while residue I4 is at proximity of desosamine sugar. TEL lacking cladinose sugar and failing to form MsrDL-SRC has been aligned (PDB 4V7S) ^79^. (j) Presence of residue I4 avoids rotation of A2062 to form an Hoogsteen base pairing with m^2^A2503 as it is the case for ErmCL-SRC (PDB 3J7Z, green)^12^. Light blue dashed lines indicate hydrogen bonds formed by Hoogsteen base pairing. For the whole figure, structures were aligned on domain V of 23S rRNA. Spheres represent van der Waals radii.

Our biochemical investigations showed that MsrDL-SRC was formed due to hampered translation termination on UAA stop codon. This stop codon is recognized by both RF1 and RF2 which subsequently catalyze peptide hydrolysis via their conserved GGQ motif protruding within the PTC. Structural alignment of our model to RF1- and RF2-containing ribosome structures^66^ revealed that proper accommodation of the RFs is prevented by steric clashes of the methylated glutamine and the following glycines of the GGQ motif with the residue F5 of MsrDL and 23S rRNA bases U2584 and U2585 respectively (Fig. 4e). The conformation of MsrDL peptide stabilizes the PTC in an uninduced conformation^67^ which prevents the opening of the active site necessary for peptide bond formation and hydrolysis (Fig. 4f). This observation suggests that MsrDL-SRC cannot catalyze peptide bond formation, as confirmed by the toeprinting preformed on the MsrDL_7A-iso_ construct (Fig. 3e). Structural alignment of MsrDL-SRC with ErmBL- and ErmCL-SRC demonstrates that 23S rRNA nucleotides U2506 and 2585 are not in the same conformation (Supplementary fig. 5a and 5b), these bases in MsrDL-SRC being in a conformation more similar to ErmDL-, SpeFL- and TnaC(R23F)-SRC (Supplementary fig. 5c to 5e)^12–16^. However, unusual position of base U2584 seems to be a unique feature of MsrDL-SRC.

MsrDL may monitor the presence of cladinose-containing drugs as our *in vivo* and *in vitro* experiments suggested. Mutation of residues L3 and I4 abolished induction of the reporter gene suggesting that they were critical for drug recognition and/or conformation of the peptide (Fig. 3d). Consistently, in our model we identified residue L3 as the closest amino acid from cladinose moiety and residue I4 as the closest from desosamine moiety (Fig. 4b and 4g).

However, similarly to ErmBL, no close contact between the nascent chain and the drug was observed^13,68^, residue L3 being ~3.8 Å away from cladinose sugar and residue I4 being ~3.9 Å away from desosamine sugar, consistently with a hydrophobic interaction between MsrDL and the drug (Fig. 4g to 4i). This observation is also corroborated by previous descriptions of pentapeptides conferring ERY resistance in which the presence of a leucine or isoleucine at position 3 was critical for drug recognition^69,70^. Moreover, the importance of 23S rRNA residue A2062 in acting as an ERY-sensor and in contributing to silence the PTC, has been demonstrated for some regulatory leader peptides^71,72^. In the case of ErmCL, the critical feature for macrolide-dependent stalling is the rotation of the base A2062 that forms an Hoogsteen base pairing with m^2^A2503^12,71,72^. The structure of MsrDL-SRC shows that residue I4 restrains rotation of base A2062 (Fig. 4j). Therefore, MsrDL induces an A2062/m^2^A2503-independent ribosome stalling^72^.

The path of MsrDL-NC within the ribosomal tunnel contrasts from all previous descriptions of elongating polypeptides (Fig. 4b and 5). Indeed, ERY-sensing ErmBL, ErmCL and ErmDL leader peptides^12–14^ engage in the NPET and skirt around ERY, while MsrDL curves before encountering the ligand then engages into a dead-end crevice at the entrance of NPET (Fig. 4b and 5b). The same observation (Fig. 5c) can be made with metabolite-sensing SpeFL/TnaC(R23F) leader peptides^15,16^ which avoid the crevice and enter the NPET. The first methionine of MsrDL-NC latches into the dead-end crevice which is delimited by 23S rRNA nucleotides U2584, U2586, G2608 and C2610 (Fig. 5a to 5c). This crevice located at the base of helix 93 (h93) is part of domain V of 23S rRNA and is conserved from bacteria to human (mitoribosome and cytosolic ribosome) (Fig. 5d, Supplementary fig. 5f and 5g). We named it proximal crevice.

**Figure 5.**
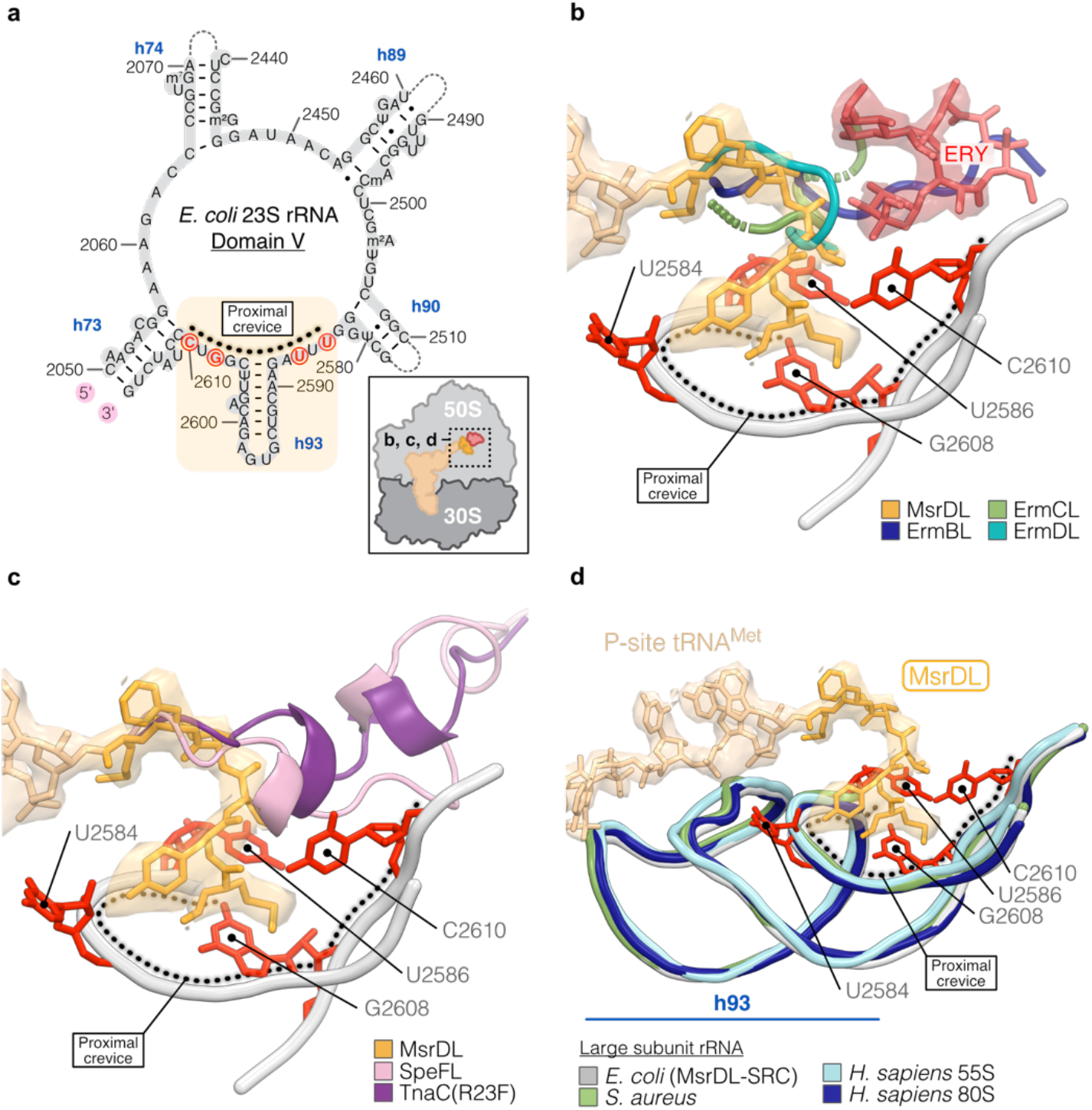
MsrDL engages in a conserved crevice at the NPET entrance. (a) Secondary structure of the *E. coli* 23S rRNA domain V showing location of the proximal crevice at the base of h93. For the whole figure, nucleotides delimitating proximal crevice are shown in red. (b and c) Comparison of MsrDL path (gold) with ERY-dependent leader peptides ErmBL (PDB 5JTE, blue), ErmCL (PDB 3J7Z, green), ErmDL (PDB 7NSO, teal) as well as L-ornithine-sensing SpeFL (PDB 6TC3, pink) and L-tryptophan-sensing TnaC(R23F) (PDB 7O1A, purple)^12–16^. See also Supplementary fig. 5a to 5e. (d) Proximal crevice in domain V is conserved from bacteria to human (MsrDL-SRC, grey; *S. aureus* PDB 6YEF, light green; *H. sapiens* 55S mitoribosome PDB 7A5F, light blue; *H. sapiens* 80S ribosome PDB 6OLI, marine blue)^80–82^. See also Supplementary fig. 5g and 5h. For the whole figure, atomic model of MsrDL is shown with its experimental density. Structures were aligned on domain V of 23S rRNA.

### MsrD negatively regulates its own expression

MsrD protein provides antibiotic resistance by rescuing ERY-stalled ribosome while *msrDL* translation in presence of ERY leads to the formation of a SRC. If MsrD is also able to rescue this SRC, it will repress its own expression and form a feedback loop similarly to Vga(A) ^73^. We tested this hypothesis using *E. coli* DB10 strains double transformed with pMMB-*msrDL-msrD*_(1-3)_:*yfp* and the compatible pBAD plasmid expressing various *msrD* variants. Strains were grown in presence of ERY at 300 nM with their optical density and fluorescence monitored (Fig. 6a). Strain carrying the *pBAD-Control* showed high fluorescence and low OD_600_ as observed in Fig. 2a, reflecting an induction of the reporter and a susceptibility to ERY. On the contrary, expression of *msrD* led to higher cell growth and a strong reduction of fluorescence which correspond respectively to antibiotic resistance and repression of the expression of the reporter. These results support that MsrD can rescue MsrDL-SRC and therefore releases the RIT repressing *msrD* transcription.

**Figure 6.**
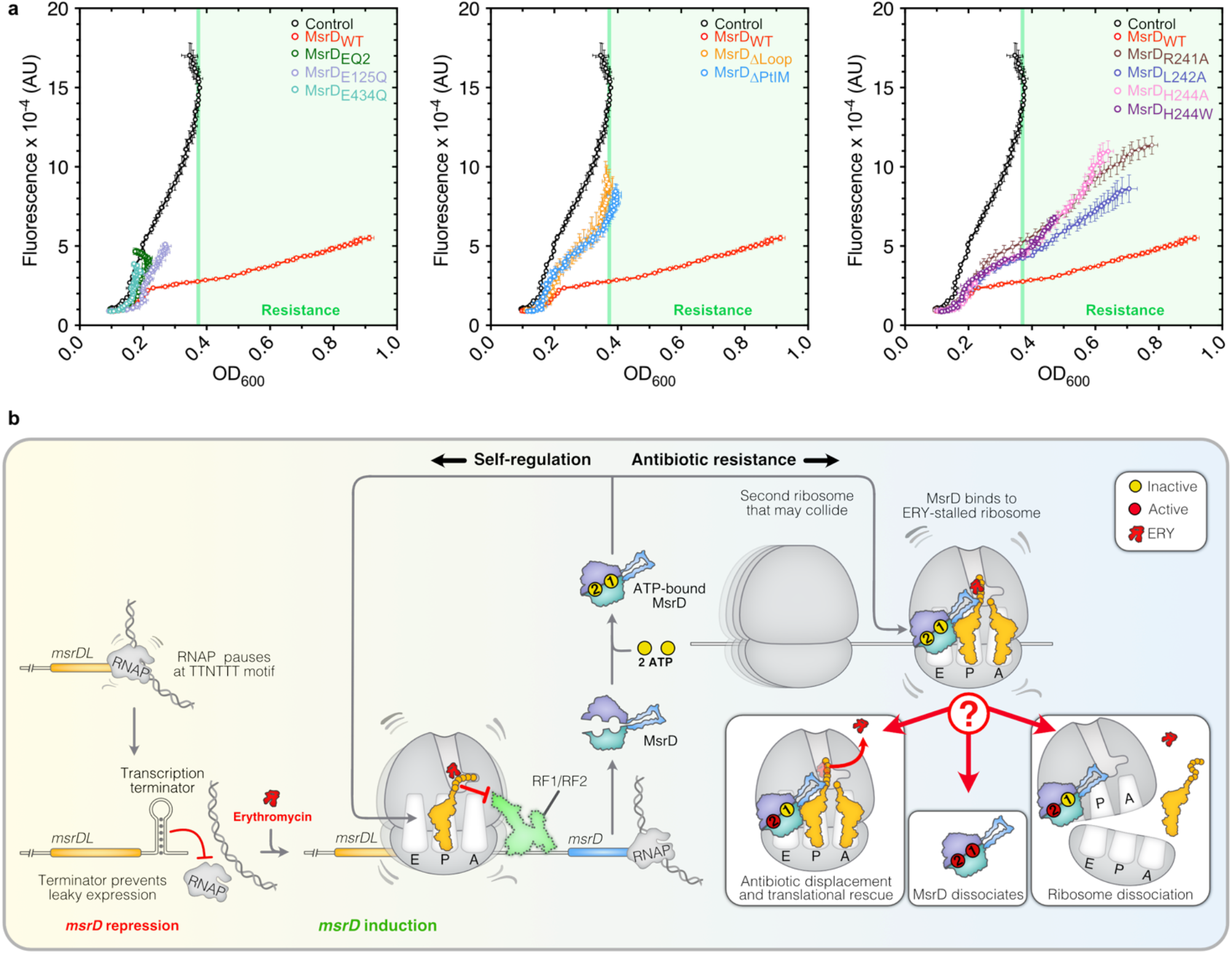
MsrD negatively regulates its own synthesis upon erythromycin exposure. (a) Effects of MsrD variants on MsrDL. *E. coli* DB10 containing *pMMB-msrDL-msrD(1-3):yfp* and expressing various *msrD* mutants were grown in presence of 0.2 % L-Arabinose, 1 mM IPTG and 300 nM ERY, both OD_600_ and fluorescence being recorded over 24 h. Fluorescence has been plotted against OD_600_, error bars for both axes represent mean ± s.d. for triplicate experiments. Color code is same as Fig. 1e to 1g. Light green rectangle indicates bacterial growth over control plasmid. (b) Model of MsrD regulating its own expression and providing antibiotic resistance. Presence of a NusG-dependent pause site may stall the RNAP and provides explanation why the system works in bacteria where transcription and translation are not so tightly coupled. In absence of ERY, RNAP drops off at Rho-independent transcription terminator. In presence of ERY, the ribosome following the RNAP (presumably paused) stalls and unwinds the terminator leading to *msrD* transcription. Once translated, MsrD negatively regulates its own expression on one side, and provides antibiotic resistance on the other side. ATP-bound MsrD recognizes ERY-stalled ribosome and may either expel the antibiotic or dissociate the ribosome, ATP site II being active. Activity in ATP site I may lead to MsrD dissociation and recycling.

The strain expressing *msrD*_E125Q_ has a low ERY resistance (Fig. 1e). When it is co-transformed with the pMMB-*msrDL-msrD*_(1-3)_:*yfp* plasmid, no resistance is observed, possibly because the presence of the two plasmid creates some toxicity. Overall, the results show that MsrD variants that lost the ability to provide antibiotic resistance (ΔPtIM, ΔLoop and ATPase deficient mutants) also lost the ability to rescue MsrDL-SRC (Fig. 6a). Point mutations in the Loop of the PtIM indicated some discrepancy between antibiotic resistance phenotype and MsrDL-SRC rescue. All the mutants show an intermediary phenotype with less antibiotic resistance and less repression of reporter compared to MsrD. The mutant MsrD_H244W_ has the same repression effect on the reporter than MsrD_H244A_ for an equivalent OD_600_ but it provides more that 3 times less antibiotic resistance (Fig. 1g and 6a). The mutant MsrD_R241A_ which provides a resistance close to MsrD does not rescue the MsrDL-SRC as efficiently as the latter one (Fig. 1g and 6a). Mutants MsrD_L242A_ and MsrD_H244A_ provide similar resistance, but MsrD_H244A_ confers a stronger repression of the reporter (Fig. 1g and 6a). Together these observations demonstrate that the determinants of the dual function of MsrD in conferring antibiotic resistance and rescuing MsrDL-SRC are overall similar.

## DISCUSSION

We used genetic, molecular biology and structural approaches to dissect the regulation of streptococcal ARE ABC-F gene *msrD*, its effect on ERY-stalled ribosomes and to provide information on ARE ABC-F mechanisms of action. We characterized the *msrDL* regulatory ORF which induces ribosome stalling in presence of cladinose-containing macrolides (Fig. 3, 4 and 6b). Our structural data show that this inhibition results from the conjoint effects of: (*i*) MsrDL maintaining 23S rRNA bases U2506 and U2585 in an uninduced conformation (Fig. 4f), (*ii*) the steric occlusion by U2584, U2585 and MsrDL residue F5 that precludes the productive accommodation of RF1/RF2 (Fig. 3c and 4e). MsrDL-stalled ribosome then hampers formation of a RIT allowing transcription of *msrD* (Fig. 2c, 3a, 3c and 6b). The coupling between translation and transcription necessary for this mechanism in *Streptococci* is possibly mediated by a NusG-dependent RNAP pausing site in *msrDL* (Fig. 2c).

MsrD binds with some specificity to ERY-stalled ribosome (Fig. 1d) and then its ATPase activity drives mechanical rearrangements to rescue stalled ribosome. The ATPase mechanism that allows the rescue is still unknown for ARE ABC-Fs and all the current structures of an ARE ABC-F in complex with the ribosome were generated with an ATPase-dead mutant^1–3,5^ or use of non-hydrolysable ATP analogues^7^. These structures do not capture an antibiotic stalled complex and embodies a reaction intermediate before or after antibiotic release of an ARE ABC-F trapped on the ribosome. We provide here some evidence about possible mechanisms (Fig. 6b). We show that only one active ATPase site (Site II, MsrD_E125Q_) can provide some antibiotic resistance (Fig. 1e) while generating a strong toxicity for the cell independently of the antibiotic and also shows more association with 50S (Fig 1c and 1d). The mutant of Site II (MsrD_E434Q_) shows a stronger toxicity (Fig. 1e) and no antibiotic resistance, effects related to less dissociation from the ribosome (Fig. 1c and 1d). Therefore, it is possible that site I is needed for complete dissociation of MsrD from the ribosome or subunits after ATP hydrolysis occurs at site II. Mutation of both sites (MsrD_EQ2_) leads to the same toxicity as for inactive Site I (Fig. 1e), suggesting that having only Site II mutated induces a new MsrD conformation that inhibits more cell growth. In accordance with these results, independent E-to-Q mutations in the ARE ABC-F Vga(A), show that mutation in site I maintains an higher ATPase activity than mutation in site II^74^.

We envision two putative non-exclusive scenarios that involve for the first one, a disassembling of ERY-stalled ribosomes, and for the other one, a displacement of the drug followed by a restart of the translation process (Fig. 6b). A key point for future studies will be to explore the ATPase activity of MsrD in a stalled-ribosome context to determine how many rounds of ATP hydrolysis are necessary to rescue the latter one. The dissociation model requires the hydrolysis of only two ATP molecules, while antibiotic displacement may require multiple hydrolysis rounds in Site II to displace the drug and to position back the elongating peptide in a correct conformation in the NPET. So far, those experiments have been challenging due to the low solubility of all the ARE ABC-F proteins^2,6^.

Alongside with ErmBL, ErmCL and ErmDL, MsrDL constitutes the fourth ERY-sensing NC structurally described, however some mechanisms of drug-sensing and PTC silencing differ. First, similarly to ErmCL, the presence of C3 cladinose sugar is required for stalling (Fig. 3a and 3b) contrary to ErmBL and ErmDL that senses drugs lacking this moiety such as oleandomycin, solithromycin or telithromycin^12,14,59,68^. Second, similarly to ErmBL, no close contact between the drug and MsrDL was observed (Fig. 4h and 4i)^68^. Third, MsrDL is the only described leader peptide that exploits inhibition of its hydrolysis by RFs to induce stalling in presence of antibiotic, similarly to the metabolite sensors TnaC and SpeFL^15,16^. On the contrary, ErmBL^13^, ErmCL^12^ and ErmDL^14^ employ only elongation inhibition. Fourth, ERY-dependent stalling on *msrDL* regulates downstream *msrD* via transcriptional attenuation (Fig. 2b and 2c) while it relies on translational attenuation for ErmBL, ErmCL and ErmDL^14,68,75^. Finally, MsrDL-NC conformation and path within the ribosome greatly differ from previously described ligand-sensing leader peptides that elongate within the tunnel (Fig. 5b and 5c)^12–16^. It adopts a hook-like shape possibly due to hydrophobic interactions with the drug (Fig. 4b, 4g and 4h), while its initiating methionine engages in a dead-end crevice at the NPET entrance (Fig. 5). To our knowledge, this is the first description of this crevice, conserved in prokaryotes and eukaryotes (Supplementary fig. 5f and 5g), as a functional site within the ribosome and we propose to name it proximal crevice.

ARE ABC-F genes tend to be transcriptionally regulated as illustrated by *msrD, vmlR, vgaA, lmo0919* and *lmrC*^52,73,76–78^. However, MsrD can regulated its own expression by creating a negative feedback-loop which is not the case for VmlR^52^. The gene *msrD* forms an operon with the efflux pump *mefA* (Fig. 1a) which is also regulated *via* ribosome-mediated transcriptional attenuation, translational stalling occurring on *mefAL*^33^. Therefore, *msrD* is under the dual-regulation of two ERY-sensing leader peptides. One explanation for this redundancy may reside in the need for the bacteria to tightly regulate *msrD* because of its toxicity. Expression of *msrD_WT_* in *E. coli* DB10 led to a 20 % growth defect compared to control condition (Fig. 1e) suggesting that *msrD_WT_* expression has a fitness cost for the bacteria and can be beneficial only in presence of antibiotic. The presence of two attenuators creates a double-lock system to avoid any basal expression in absence of inducer and the negative feedback-loop maintains a minimal amount of MsrD production. Thus, as long as MefA can export enough antibiotic to provide resistance, *msrD* should be kept repressed. The fitness cost of the expression of ARE ABC-F genes is possibly the Achilles heel of bacteria containing those genes and may be exploited to fight antibiotic resistance. MsrD provides resistance to ERY, AZY and TEL^23^ (Table 1) but only the first two induce its expression, indicating that MsrD can provide some resistance for an antibiotic which does not induce its expression. Thus, MsrDL facilitates the use of MsrD by the bacteria and maintains its evolution under its control.

## Supporting information

SI_Fostier_et_al_Aug_2022

## ACKNOWLEDGMENTS

This work has benefited from the facilities and expertise of the Biophysical and Structural Chemistry platform (BPCS) at IECB, CNRS UMS3033, Inserm US001, University of Bordeaux. C.R.F. is funded by a doctoral grant from the french Ministère de l’Enseignement supérieur, de la Recherche et de l’Innovation, F.O. and G.B. have received support from the LABEX program (DYNAMO ANR-11-LABX-0011), and the ANR grants EZOtrad (ANR-14-ACHN-0027) for G.B. and ABC-F_AB (ANR-18-CE35-0010) for G.B. and Y.H., ERC-2017-STG #759120 “TransTryp” to Y.H. E.C.L. and C.A.I. have received funding for this project from the European Research Council (ERC) under the European Union’s Horizon 2020 research and innovation program (Grant Agreement No. 724040). Authors would like to thank Dr. Olivier Chesneau (Département de Microbiologie, Institut Pasteur, Paris - France) for providing *E. coli* DB10 strain, plasmids pBAD33 and pVN50; Dr. Sylvain Durand and Dr. Maud Guillier (UMR 8261, CNRS, Université de Paris, Institut de Biologie Physico-Chimique, Paris - France) that kindly provided respectively bicyclomycin and equipment for toeprinting experiments. Authors would like also to thank Laura Monlezun and Tina wang for proofreading the manuscript.

## AUTHOR CONTRIBUTIONS

F.O. performed the first experiments that initiated the project. C.R.F. performed most of the experiments. S.N. performed northern blotting experiments. C.R.F. prepared the cryo-EM sample. H.S. prepared cryo-EM grids and imaged them. C.A.I. and Y.H. processed cryo-EM data. C.R.F and E.C.L. reconstructed the atomic model. G.B. initiated the project and designed the research program with Y.H. C.R.F. and G.B. wrote the paper with input from all authors.

## DECLARATION OF INTERESTS

The authors declare no competing interests. Requests for materials should be addressed to G.B.

## METHOD DETAILS

### Construction of plasmids

Strains and plasmids used in this study are listed in Supplementary Table 3. Oligonucleotides are listed in Supplementary Table 4. For the whole study, the considered reference sequence for *mefA/msrD* macrolide resistance operon was a Tn916-type transposon inserted in *Streptococcus pneumoniae* strain 23771 genome (Genbank accession number FR671415.1)^83^. Plasmids *pBAD-Control* and pBAD-*msrD*_WT_ containing a C-terminal hexahistidine tag (originally referred as pBAD33 and pVN50) were kindly retrieved from Olivier Chesneau at Institut Pasteur (Paris, France)^24^. Both *mefA/msrD* macrolide resistance operon and pBAD-*msrD*_WT_ encode the same protein MsrD (Uniprot accession number A0A496M710).

#### pBAD plasmids

Those plasmids allow a tight and stringent expression control via catabolite repression, interest gene being repressed in presence of 0.4 % (w/v) β-D-Glucose and induced in presence of 0.2 % (w/v) L-Arabinose. Transformed clones were selected with 20 μg.ml^-1^ chloramphenicol. In this study, all *msrD* mutants contain a C-terminal hexahistidine tag. Catalytic mutant *msrD*_EQ2_ was generated by amplifying pBAD-*msrD*_WT_ with primer pairs 5/6 and 7/8, both fragments being assembled with NEBuilder HiFi DNA Assembly Master Mix (New England Biolabs). Catalytic mutants *msrD*_E125Q_, *msrD*_E434Q_ were generated via quickchange mutagenesis with primers pairs 9/10 and 11/12. Mutants *msrD*_ΔLoop_ and *msrD_ΔPtlM_* were generated based on phylogenetic alignments by deleting residues between K216/K254 and E189/A274 respectively, a three-glycine linker being added to allow flexibility, via quickchange mutagenesis using primer pairs 13/14 and 15/16. Variants *msrD*_R241A_, *msrD*_L242A_, *msrD*_H244A_, *msrD*_H244W_ were generated via fusion PCR by amplifying *MsrD_WT_* with primers pairs 1/18, 1/20, 1/22, 1/24 and 2/17, 2/19, 2/21, 2/23 respectively. Backbone was amplified with primer pair 3/4 and fragments were assembled with NEBuilder HiFi DNA Assembly Master Mix (New England Biolabs).

#### pMMB plasmids

All pMMB constructs originated from a low copy IPTG-inducible plasmid pMMB67EH-*yfp*, consisting in original pMMB67EH^45^ containing optimized *venus-yfp* enhancing its translation under the control of P_tac_ promoter (manuscript in preparation). Transformed clones were selected with 100 μg.ml^-1^ ampicillin. To test *msrDL* functions *in vivo*, several fluorescent reporter genes have been designed. First, pMMB67EH-*yfp* native P_tac_ promoter was replaced by P_LlacO-1_ promoter as previously described^46^: 5’-ATAAATGTGAGCGGATAACATTGACATTGTGAGCGGATAACAAGATACTGAGCACA-3’ (lac operators, shaded grey; transcription start, bold). This IPTG-inducible promoter allows a tight and stringent transcription regulation compared to P_tac_ promoter, insofar as no regulatory element is found in the 5’ UTR. Promoter was replaced by amplifying pMMB67EH-*yfp* with primer pairs 25/26, PCR fragment being then re-circularized via NEBuilder HiFi DNA Assembly

Master Mix (New England Biolabs). The resulting plasmid was named pMMBpLlacO-1-67EH*-yfp.* Plasmid *pMMB-msrDL-msrD_(1-3)_:yfp* has been designed by introducing the sequence spanning from the first nucleotide downstream *mefA* stop codon to *msrD* three first codons fused to *yfp*. The introduced sequence is as follows: 5’-ACAATATT**GGAGGA**ATATTTATGTATCTTATTTTCATGTAACTCTTCCTGCTAAAATCGCA GGGTTTTCCCTGCATACAAGCAAATGAAAGCATGCGATTATAGACA**GGAGGA**AATGTTA TGGAATTA-3’ (RBS, bold; *msrDL*, shaded grey, *msrD* three first codons are underlined). To clone such construct, *yfp* gene was amplified from pMMB67EH-*yfp* with primer pairs 27/29 then with 28/29 to generate the insert, backbone was amplified with primer pair 30/31 using pMMBpLlacO-1-67EH*-yfp* as matrix, both fragments being then assembled with NEBuilder HiFi DNA Assembly Master Mix (New England Biolabs). Plasmid pMMB-*control* was built by removing the *msrDL*-*msrD*_(1-3)_:*yfp* cassette via quickchange mutagenesis using primer pair 32/33, the resulting construction containing only the P_LlacO-1_ promoter followed by the plasmid endogenous *rrnB* transcription terminator. Plasmid pMMB-*msrDL*_(no_term)_ *-msrD*_(1-3)_*:yfp* (where the RIT between *msrDL* and *msrD* is deleted) was generated by amplifying pMMB-*msrDL*-*msrD*_(1-3)_:*yfp* with primer pair 34/35, PCR fragment being then re-circularized via NEBuilder HiFi DNA Assembly Master Mix (New England Biolabs). The various *msrDL* mutants were cloned via quickchange mutagenesis by amplifying pMMB-*msrDL*-*msrD*_(1-3)_*:yfp* with primers 36 to 51. Plasmid pMMB-*msrDL*_(MYLIFMA-isocodons)_-*msrD*_(1-3)_:*yfp* was generated by replacing *msrDL*_WT_ sequence (ATGTATCTTATTTTCATGTAA) by a recoded sequence (ATGTACCTGATCTTCATGGCCTAA) using isocodons and introducing an extra alanine codon (7A codon) before stop codon. Sequence was recoded using isocodons because mutating the WT sequence introduced a new promoter. To do so, pMMB-*msrDL*-*msrD*_(1-3)_*:yfp* was amplified with primer pair 52/53, PCR fragment being then re-circularized via NEBuilder HiFi DNA Assembly Master Mix (New England Biolabs). Recoded sequence without the 7A codon (ATGTACCTGATCTTCATGTAA) was generated via quickchange mutagenesis by amplifying pMMB-*msrDL*_(MYLIFMA-isocodons)_-*msrD*_(1-3)_:*yfp* with primer pair 54/55, leading to plasmid pMMB-*msrDL*_(WT-isocodons)_*-msrD*_(1-3)_:*yfp.*

### Antibiotic susceptibility testing, MIC and IC50 determination

A saturated preculture of *E. coli* DB10 transformed with pBAD plasmid was grown overnight at 37 °C under vigorous shaking in Luria-Bertani Miller broth (LB), 20 μg.ml^-1^ chloramphenicol and supplemented with 0.4 % (w/v) β-D-Glucose. Antibiotic susceptibility testing assay was performed in Mueller-Hinton broth (MH, Sigma Aldrich), antibiotics being diluted via serial dilutions. A 96-wells flat-bottom plate (Flacon) was filled with a final volume per well of 200 μl, containing 20 μg.ml^-1^ chloramphenicol and 0.2 % (w/v) L-Arabinose and antibiotic to test. Wells were inoculated at OD_600_ ~0.03-0.04 prior to addition of 60 μl mineral oil (Sigma Aldrich) avoiding evaporation but not oxygen diffusion. Plates were therefore incubated for 24 h in CLARIOstar Plus plate reader (BMG Labtech) at 37 °C with 600 rpm double-orbital shaking, OD_600_ being measured each 30 min. Optical densities at 24 h were then normalized relative to optical densities of *E. coli* DB10 *pBAD-Control* grown in the absence of antibiotic in Prism 7(GraphPad). MIC was determined as absence of growth compared to blank, IC50 was calculated in Prism using equation Y=Bottom + (Top-Bottom)/(1 + ((X^HillSlope)/(IC50^HillSlope))) and the standard deviation was calculated by multiplying standard error by square root of n (n being at least 3 replicates). To generate curves, for bacteria grown in absence of antibiotic (0 μM), since coordinates are plotted as logarithms and since log(0) is undefined, this point has been approximated 2 log units below the lowest tested value *(i.e.* 0.0625 μM) consistently with Prism user guide^84^. Curve fitting was performed with non-linear fitting function “log(inhibitor) vs. response -- Variable slope” using Y=Bottom + (Top-Bottom)/(1+10^((LogIC50-X)*HillSlope)). Both equations gave strictly the same result for IC_50_.

### *In vivo* induction assay

A saturated preculture of *E. coli* DB10 transformed with pMMB plasmid was grown overnight at 37 °C under vigorous shaking in LB with 100 μg.ml^-1^ ampicillin. For bacteria double-transformed with pMMB and pBAD plasmids, media was also supplemented with 20 μg.ml^-1^ chloramphenicol and supplemented with 0.4 % (w/v) β-D-Glucose. *In vivo* induction assay was performed in Mueller-Hinton broth (MH, Sigma Aldrich). A 96-wells flat-bottom plate (Flacon) was filled with a final volume per well of 200 μl, containing 100 μg.ml^-1^ ampicillin, 1 mM Isopropyl β-D-1-thiogalactopyranoside (IPTG). Growth medium was supplemented with 20 μg.ml^-1^ chloramphenicol and 0.2 % (w/v) L-Arabinose if pBAD plasmid is present. Induction of fluorescent reporters being antibiotic-dependent, growth media were supplemented with required antibiotic accordingly. Plates were therefore incubated in CLARIOstar Plus plate reader (BMG labtech) at 37 °C with 600 rpm double-orbital shaking up to 24 h, OD_600_ and fluorescence (excitation: 497-15 nm, emission: 540-20 nm, gain 1600) being measured each 30 min.

### Polyribosomes fractionation

Saturated precultures of *E. coli* DB10 transformed with pBAD-*control*, pBAD-*msrD*_WT_, pBAD-*msrD*_EQ2_, pBAD-*msrD*_E125Q_ or pBAD-*msrD*_E434Q_ were grown overnight at 37 °C under vigorous shaking in LB, 20 μg.ml^-1^ chloramphenicol, 0.4 % (w/v) β-D-Glucose. Growth media (MH, 20 μg.ml^-1^ chloramphenicol, 0.2 % (w/v) L-Arabinose) was inoculated at OD_600_=0.05 with normalized precultures, and cells were grown at 37 °C and 180 rpm. When OD_600_=0.5 was reached, cultures were divided in two and one half was treated with a 25 μM erythromycin.

Cultures were then incubated 1 hour at 37 °C and 180 rpm before harvesting. Cultures were chilled and cells were harvested by centrifuging 10 min at 8 000 rpm, 4 °C. Supernatant was discarded and cells were resuspended in 350 μl of lysis buffer (20 mM Tris pH 8, 20 mM MgCl_2_, 100 mM NH_4_(OAc), 2 mM β-mercaptoethanol, 1 mg.ml^-1^ lysozyme) and transferred in microtubes. Then, 30 μl of sodium deoxycholate 10 % (w/v) was added and cells were mechanically lysed by 3 rounds of freezing-thawing. Lysates were clarified by centrifuging 20 min at 18 000 xg, 4 °C and total RNAs were quantified. Finally, 500 μg of total RNAs were loaded onto 10-40 % (w/v) sucrose gradients (20 mM Tris pH 8, 20 mM MgCl_2_, 100 mM NH_4_(OAc), 2 mM β-mercaptoethanol) and centrifuged 2 h at 40 000 rpm, 4 °C in SW41 rotor (Beckman Coulter). Samples were fractionated and collected using Biocomp Piston Gradient Fractionator (BioComp Instruments), absorbance being measured at 254 nm.

### Immunoblotting

Fractions of 400 μl were precipitated in ice-cold absolute ethanol (1 volume of sample, 3 volumes of ethanol) overnight at −20 °C. Samples were then centrifuged 45 min at 18 000 xg, 4 °C. Supernatant was removed and samples were vacuum-dried. Pellets were resuspended in 33.3 μl 1X Laemmli buffer, 10 μl being loaded on 12.5 % acrylamide SDS-PAGE gels. Concerning total lysates, 10 μg of normalized total RNAs were loaded as control. Gels were resolved by migrating at 0.04 A and then applied on Immun-Blot PVDF membrane (Bio-Rad) that had been activated in absolute ethanol then washed in electro-transfer buffer (25 mM Tris base, 192 mM glycine, 0.1 % (w/v) SDS, 10 % (v/v) absolute ethanol). Proteins were electro-transferred at 100 V during 1 h. Membranes were blocked by incubating 1 h in 1X PBS supplemented with 0.5 % nonfat dry milk then washed with 1X PBS. C-term His-tagged MsrD_WT_ and MsrD_EQ2_ were detected using anti-6xHis-tag primary antibody (Covalab, 1:2 000 dilution in 1X PBS, 0.1 % Tween-20) combined with anti-mouse-HRP secondary antibody (Covalab, 1:20 000 dilution in 1X PBS, 0.1 % Tween-20). Immunoblots were revealed by performing an ECL detection using Clarity Western ECL Substrate (Bio-Rad) and imaged with Li-Cor Odyssey FC imaging system (Li-Cor).

### Preparation of *E. coli* DB10 ribosomes

An overnight saturated *E. coli* DB10 preculture was used to inoculate growth medium (LB, dilution 1:100) and cells were grown at 37 °C and 180 rpm. When OD_600_=0.5, cells were washed and harvested by two centrifugations during 20 min, 5 000 xg at 4°C followed by resuspensions in Buffer A (20 mM Tris pH 7.4, 10 mM Mg(OAc), 100 mM NH_4_(OAc), 0.5 mM EDTA). Pellets were resuspended in Buffer A supplemented with 6 mM β-mercaptoethanol, 10 mg.ml^-1^ lysozyme, 0.001 % (v/v) protease inhibitor cocktail (Sigma Aldrich) and lysed three times at 2.5 kBar using a cell disrupter (Constant Systems Limited). Lysate was clarified by two centrifugations during 15 min, 22 000 xg at 4°C then spun for 20 h, 34 700 rpm at 4 °C in a Type 70 Ti rotor (Beckmann Coulter) through a 37.7 % (w/v) sucrose cushion in Buffer B (20 mM Tris pH 7.4, 10 mM Mg(OAc), 500 mM NH4(OAc), 0.5 mM EDTA, 6 mM β-mercaptoethanol). Sucrose cushions were decanted and ribosome pellets resuspended in Buffer C (20 mM Tris pH 7.4, 7.5 mM Mg(OAc), 60 mM NH4(OAc), 0.5 mM EDTA, 6 mM β-mercaptoethanol). Finally, 12 mg of ribosomes were loaded onto 10-40 % (w/v) sucrose gradients in Buffer C and centrifuged 18 h at 20 000 rpm, 4 °C in SW28 rotor (Beckman Coulter). Gradients were fractionated using Biocomp Piston Gradient Fractionator (BioComp Instruments), absorbance being measured at 254 nm. Fractions containing 70S ribosomes were pooled, washed and concentrated in Amicon 50k (Merckmillipore) using Buffer C. Ribosomes concentration was adjusted to 18-20 μM, then aliquoted and flash frozen in liquid nitrogen.

### RNA extraction and northern blotting

Total RNA extraction was realized using the RNAsnap method as previously described^85^. In brief, a preculture of *E. coli* DB10 containing *pMMB-msrDL-msrD_(1-3)_:yfp* was grown overnight at 37 °C under vigorous shaking in LB supplemented with 100 μg.ml^-1^ ampicillin. Growth medium (MH, 100 μg.ml^-1^ ampicillin) was inoculated at OD6_00_=0.05, and cells were grown at 37 °C and 180 rpm. When OD_600_=0.5, cells were treated (or not) with 1 mM IPTG during 10 min, then treated (or not) with 100 nM erythromycin. Samples of 2 ml were collected (T_0’_ = before IPTG was added, T_10’_= 10 min after IPTG was added, and 5 min, 15 min, 30 min, 60 min and 120 min after erythromycin was added), then spun during 1 min at 15 000 rpm. Growth media was removed and cells were resuspended in 100 μl RNAsnap buffer (95 % (v/v) formamide, 18 mM EDTA, 0.025 % (v/v) SDS, 1 % (v/v) β-mercaptoethanol). Samples were heated for 10 min at 95 °C then clarified by centrifuging 10 min, at 16 000 rpm at 16 °C. For northern blotting analysis, 6 μg of total RNAs were resolved on a 1 % (w/v) agarose gel then transferred onto Amersham Hybond-N+ Membrane (GE Healthcare) by capillary transfer. Radioactive probe was prepared using 40 pmol of primer 59 and 5’-labelled with 10 U of T4 Polynucleotide Kinase (New England Biolabs) and [γ^32^P]ATP (150 μCi). Probe was hybridized overnight at 42 °C using ULTRAhyb-Oligo hybridization buffer (Thermo Fisher Scientific). Membrane was washed three times at 42 °C during 15 min (once in 2x SSC + 0.1 % (v/v) SDS, once in 1x SSC + 0.1 % (v/v) SDS and finally in 0.1x SSC + 0.1 % (v/v) SDS). Radioactive signal was visualized by exposing 4 h a Storage Phosphor Screen BAS-IP MS 2040 (Fujifilm) then imaged with a Typhoon FLA 9500 (GE Healthcare).

### *In vitro* transcription

*In vitro* transcription was carried out in T7 RiboMAX Large Scale RNA Production System kit (Promega) according to manufacturer instructions. Briefly, to generate DNA matrix (Supplementary Table 5), T7 promoter (5’-GCGAATTAATACGACTCACTATAGGG-3’) was added by PCR using primers pair 56/57 (Supplementary Table 4) and pMMB plasmids as templates. Transcription reactions were incubated 4 h at 37 °C and transcripts were purified with TRIzol Reagent (Thermo Fisher Scientific) and Direct-zol RNA Miniprep kit (Zymo Research), samples being eluted in THE Ambion RNA Storage Solution (Thermo Fisher Scientific). Final concentration was adjusted to 5 pmol.μl^-1^.

### Toeprinting assay

Position of stalled ribosomes on mRNA was determined by toeprinting assay, slightly adapted from previously described methods^15,86^. *In vitro* translation reactions were performed using PURExpress ΔRF123 kit and PURExpress ΔRibosome (New England Biolabs) according to manufacturer instructions. Briefly, prior to *in vitro* translation reactions, 0.5 μl of 500 μM ligand (e.g. erythromycin etc) was dried in a micro-centrifuge tube, using a SpeedVac vacuum concentrator (Thermo Fisher Scientific), final concentration being 50 μM once resuspended in a 5 μl reaction. Reactions were incubated during 15 min at 37 °C using 5 pmol of RNA templates (generated as described above) and 3.3 μM of purified *E. coli* DB10 ribosomes. When needed, samples were treated with 100 μM final puromycin and incubated 3 min more. Immediately, 2 pmol of CY5-labelled primer 58 complementary to NV1 sequence ^20^ was added and incubated for 5 min at 37 °C. For each reaction, reverse transcription was performed with 0.5 μl (corresponding to 5 U) of Avian Myeloblastosis Virus Reverse Transcriptase (Promega), 0.1 μl dNTP mix (10 mM), 0.4 μl Pure System Buffer (5 mM K-phosphate pH 7.3, 9 mM Mg(OAc), 95 mM K-glutamate, 5 mM NH4Cl, 0.5 mM CaCl2, 1 mM spermidine, 8 mM putrescine, 1 mM DTT) and incubated 20 min at 37 °C. Reactions were quenched with 1 μl NaOH 5 M and incubated 15 min at 37 °C. Alkali were therefore neutralized by adding 0.7 μl HCl 25% immediately supplemented with 20 μl toe-printing resuspension buffer (300 mM Na(OAc) pH 5.5, 5 mM EDTA, 0.5% (v/v) SDS). Finally, cDNAs were purified using QIAquick Nucleotide Removal kit (Qiagen), vacuum dried and resuspended in 6 μl formamide dye (95% (v/v) formamide, 20 mM EDTA, 0.25% (w/v) bromophenol blue). Sanger sequencing was performed on DNA matrix used for *in vitro* transcription. Briefly, each 20 μl reaction contained 7.5 nM DNA matrix, 75 nM CY5-labelled primer 58, 40 μM of each dNTPs, 0.025 U Taq Pol (New England Biolabds), 1x Thermo Pol Buffer (New England Biolabs) and corresponding ddNTPs (625 μM ddCTP/ddTTP/ddATP or 50 μM ddGTP). After PCR, 20 μl formamide dye was added to each sequencing reaction. Samples were denatured 3 min at 80 °C, then 5 μl of sequencing reaction and 1.5 μl of toe-printing reaction were loaded on a 6% acrylamide/bis-acrylamide (19:1) sequencing gel containing 8 M urea. Gel was resolved by migrating 90 min at 50 W, then imaged using a Typhoon FLA 9500 (GE Healthcare Life Sciences) using CY5 mode, LPR Ch.2 filter and 635 nm laser.

### Preparation of MsrDL-SRC

A DNA matrix (Supplementary Table 5) was prepared with CloneAmp HiFi PCR Premix (Takara) using primer pair 56/60 (Supplementary Table 4) and plasmid pMMB-*msrDL*-*msrD(1-3):yfp* as matrix. The corresponding mRNA was generated as described above. The complete sequence of MsrDL-SRC mRNA is: 5’-GGAGCGGAUAACAAGAUACUGAGCACAACAAUAUU**GGAGG**AAUAUUUAUGUAU CUUAUUUUC<u>AUG</u>UAACUCUUCCUGCUAAAAUCGCAGGGUUUUCCCUGC-3’ (RBS, bold; *msrDL,* shaded grey; the ATG codon in the P site of stalled ribosomes is underlined). Prior to *in vitro* translation reaction, 1 μl of 500 μM erythromycin (50 μM final concentration once resuspended in 10 μl) has been dried in a micro-centrifuge tube, using a SpeedVac vacuum concentrator (Thermo Fisher Scientific). Dried erythromycin has been resuspended in a 10 μl *in vitro* translation reaction carried out in PURE*frex* 2.0 kit (Genefrontier), containing 1.83 μM of purified *E. coli* DB10 ribosomes and 3.6 μM mRNA (molar ratio 1:2). The reaction has been incubated for 10 minutes at 37 °C, before adding 100 μM final puromycin, and incubated 3 min more. Reaction was therefore diluted in ribosome purification Buffer C (20 mM Tris pH 7.4, 7.5 mM Mg(OAc), 60 mM NH4(OAc), 0.5 mM EDTA, 6 mM β-mercaptoethanol) to reach a concentration of 150 nM ribosomes. Cryo-EM grids were immediately prepared.

### Cryo-EM grids preparation

Safematic ccu-010 HV carbon coater was used to coat Quantifoil carbon grids (QF-R2/2-Cu) with a thin carbon layer of approximate thickness of 2nm. Grids were therefore glow discharged for 30 sec at 2 mA. Then, 4 μl of *in vitro* translation reaction diluted to 150 nM were applied, and after a 2 sec blotting (force 5) and 30 sec waiting time, grids were vitrified in liquid ethane using a Vitrobot Mark IV (FEI) set to 4 °C and 100 % humidity.

### Image acquisition and processing

Data collection was performed on a Talos Arctica instrument (FEI Company) at 200 kV using the EPU software (Thermo Fisher Company) for automated data acquisition. Data were collected at a nominal defocus of −0.5 to −2.7 μm at a magnification of 120,000 X yielding a calibrated pixel size of 1.2 Å. Micrographs were recorded as movie stack on a K2 Summit direct electron detector (Gatan), each movie stack were fractionated into 65 frames for a total exposure of 6.5 sec corresponding to an electron dose of 64 e^-^/Å^2^. MotionCor2 ^87^ was used for dose weighting, drift and whole-frame motion correction. A dose weighted average image of the whole stack was used to determine the contrast transfer function with the software Gctf^88^. Particles were picked using a Laplacian of gaussian function (min diameter 260 Å, max diameter 320 Å). A total of ~254k particles were extracted from a subset of motion-corrected images (1523 micrographs) presenting a resolution equal or better than 4 Å with a box size of 360 pixels. The particles were binned three folds for 2D and subsequent 3D classification. After 2 rounds of 2D classification in RELION 3^89^, ~158k particles were selected and submitted to Relion 3D classification^89^. A class of non-rotated 70S depicting high-resolution features and bearing a tRNA with various occupancies was selected, representing ~104k particles. This class was further 3D-classified with a spherical mask engulfing the tRNAs binding sites (Supplementary Fig. 4b) into 4 classes, thus yielding to P- and E-tRNAs, P-tRNAs only, A- and P-tRNAs and E-tRNAs 70S reconstructions. Only P- and E-tRNAs and P-tRNAs only reconstructions were used for further processing, which represents ~62k particles displaying the same global conformation of the 70S. These particles where re-extracted at the full pixel size (1.2 Å) and refined through the 3D auto-refinement performed in RELION 3^89^ resulting in a 3 Å reconstruction (Supplementary Fig. 4c), after CTF-refinement, Bayesian particle polishing and post-processing in RELION 3. In spite of the sufficient local resolution at the vicinity of MsrDL peptide at the PTC (~3Å), residual movements of the 30S around the 5OS can be deduced by the lower local resolutions of the head and the body of the 30S. In order to improve their resolution so to derive a complete atomic model of the entire 70S complex, we applied RELION 3 multi-body refinement by defining three bodies; 50S, 30S-head and 30S-body. After completion of the refinement, the reconstructions of the 30S-head, 30S-body and 50S reached average resolutions of 3.3, 3.08 and 2.97 Å, respectively. The local resolution (Supplementary Fig. 4b and 4d) was estimated using ResMap^90^.

### Model building and refinement

The atomic model of erythromycin-stalled *Escherichia coli* 70S ribosome with streptococcal MsrDL nascent chain was built into cryo-EM maps using Coot and Phenix^91,92^. Insofar as *E. coli* DB10 hasn’t been sequenced yet, we assumed that ribosomal proteins and rRNAs were strictly identical to *E. coli* K12. Furthermore, we did not notice any significant features in the map. Structure of SpeFL-SRC in response to L-ornithine was used as initial model (PDB 6TC3)^15^ and has been fitted in 70S ribosome map EMD-13805. Then, each part of the ribosome (50S, 30S Body and 30S Head) has been individually inspected in the corresponding map (respectively EMD-13806, EMD-13807 and EMD-13808) modified if necessary in Coot. P-site elongator tRNA _Met_tRNA^Met^ was modeled *de novo* based on *E. coli* K12 MG1655 *metT* gene, posttranscriptional modifications were added consistently with Modomics database^93^ (http://genesilico.pl/modomics/). E-site tRNA _Phe_tRNA^Phe^ was derived from crystal structure of phenylalanine tRNA from *E. coli* (PDB 6Y3G) with minor adjustments^94^. ERY, MsrDL leader peptide and mRNA were modeled *de novo* in Coot. Final model was refined in map EMD-13805 using Phenix^92^.

### Figure preparation

Growth curves, histograms and polyribosomes profiles were generated using GraphPad Prism 7 (GraphPad). Sequence alignments were visualized with JalView^95^. Western blotting, northern blotting and toe-printing gels were analyzed using Fiji^96^. Figures depicting molecular structures or electronic density maps were prepared using PyMOL Molecular Graphics System, Chimera and ChimeraX^97^,^98^.

### Quantification and statistical analysis

Statistical details can be found in the figure legends. Statistical significance was assessed using unpaired t-Test function in Prism 7 (GraphPad).

### Data and software availability

Cryo-EM map of erythromycin-stalled *Escherichia coli* 70S ribosome with streptococcal MsrDL nascent chain has been deposited at the Electron Microscopy Data Bank (EMDB) with accession code EMD-13805, as well as 50S, 30S Body and 30S Head maps obtained after multibody refinement with accession code EMD-13806, EMD-13807, EMD-13808 respectively. Corresponding atomic model has been deposited in the Protein Data Bank (PDB) with accession code 7Q4K.

## Notes

### Competing Interest Statement

The authors have declared no competing interest.

### Summary of Updates

Updated version of the manuscript. Title has been simplified. Abstract has been updated in view of new results. Main text has been modified in view of new results. Materials and methods have been updated accordingly. Figures 1C and 1D are new results. Previous Figures 1C to 1E are now in supplementary informations. Figure 2C has been modified by adding NusG pause motif. Figure 3D has been slightly modified by adding name and amino acid sequences of corresponding mutants. Current Figure 3E is a new result. Previous Figure 3E is now in supplementary informations. Figure 6B has been slightly modified to update general model. Supplementary Figure 4 has been corrected.

## REFERENCES

1. Crowe-McAuliffe, C. et al. Structural basis for antibiotic resistance mediated by the Bacillus subtilis ABCF ATPase VmlR. PNAS 115, 8978–8983 (2018).

2. Crowe-McAuliffe, C. et al. Structural basis of ABCF-mediated resistance to pleuromutilin, lincosamide, and streptogramin A antibiotics in Gram-positive pathogens. Nat Commun 12, 3577 (2021).

3. Crowe-McAuliffe, C. et al. Structural basis for PoxtA-mediated resistance to phenicol and oxazolidinone antibiotics. Nat Commun 13, 1860 (2022).

4. Lenart, J., Vimberg, V., Vesela, L., Janata, J. & Balikova Novotna, G. Detailed Mutational Analysis of Vga(A) Interdomain Linker: Implication for Antibiotic Resistance Specificity and Mechanism. Antimicrob Agents Chemother 59, 1360–1364 (2015).

5. Mohamad, M. et al. Sal-type ABC-F proteins: intrinsic and common mediators of pleuromutilin resistance by target protection in staphylococci. Nucleic Acids Research 50, 2128–2142 (2022).

6. Sharkey, L. K. R., Edwards, T. A. & O’Neill, A. J. ABC-F Proteins Mediate Antibiotic Resistance through Ribosomal Protection. mBio 7, (2016).

7. Su, W. et al. Ribosome protection by antibiotic resistance ATP-binding cassette protein. PNAS 115, 5157–5162 (2018).

8. Boёl, G. et al. The ABC-F protein EttA gates ribosome entry into the translation elongation cycle. Nat Struct Mol Biol 21, 143–151 (2014).

9. Chen, B. et al. EttA regulates translation by binding the ribosomal E site and restricting ribosome-tRNA dynamics. Nat Struct Mol Biol 21, 152–159 (2014).

10. Fostier, C. R. et al. ABC-F translation factors: from antibiotic resistance to immune response. FEBS Letters 595, 675–706 (2021).

11. Ousalem, F., Singh, S., Chesneau, O., Hunt, J. F. & Boël, G. ABC-F proteins in mRNA translation and antibiotic resistance. Research in Microbiology 170, 435–447 (2019).

12. Arenz, S. et al. Drug Sensing by the Ribosome Induces Translational Arrest via Active Site Perturbation. Molecular Cell 56, 446–452 (2014).

13. Arenz, S. et al. A combined cryo-EM and molecular dynamics approach reveals the mechanism of ErmBL-mediated translation arrest. Nat Commun 7, (2016).

14. Beckert, B. et al. Structural and mechanistic basis for translation inhibition by macrolide and ketolide antibiotics. Nat Commun 12, 4466 (2021).

15. Herrero del Valle, A. et al. Ornithine capture by a translating ribosome controls bacterial polyamine synthesis. Nature Microbiology 5, 554–561 (2020).

16. van der Stel, A.-X. et al. Structural basis for the tryptophan sensitivity of TnaC-mediated ribosome stalling. Nat Commun 12, 5340 (2021).

17. Horinouchi, S. & Weisblum, B. Posttranscriptional modification of mRNA conformation: Mechanism that regulates erythromycin-induced resistance. PNAS 77, 7079–7083 (1980).

18. Ito, K. & Chiba, S. Arrest Peptides: Cis-Acting Modulators of Translation. Annu. Rev. Biochem. 82, 171–202 (2013).

19. Narayanan, C. S. & Dubnau, D. Evidence for the translational attenuation model: ribosome-binding studies and structural analysis with an in vitro run-off transcript of ermC. Nucleic Acids Res 13, 7307–7326 (1985).

20. Vazquez-Laslop, N., Thum, C. & Mankin, A. S. Molecular Mechanism of Drug-Dependent Ribosome Stalling. Molecular Cell 30, 190–202 (2008).

21. Gay, K. & Stephens, D. S. Structure and Dissemination of a Chromosomal Insertion Element Encoding Macrolide Efflux in Streptococcus pneumoniae. The Journal of Infectious Diseases 184, 56–65 (2001).

22. Roberts, M. C. et al. Nomenclature for Macrolide and Macrolide-Lincosamide-Streptogramin B Resistance Determinants. Antimicrob Agents Chemother 43, 2823–2830 (1999).

23. Daly, M. M., Doktor, S., Flamm, R. & Shortridge, D. Characterization and Prevalence of MefA, MefE, and the Associated msr(D) Gene in Streptococcus pneumoniae Clinical Isolates. J Clin Microbiol 42, 3570–3574 (2004).

24. Nunez-Samudio, V. & Chesneau, O. Functional interplay between the ATP binding cassette Msr(D) protein and the membrane facilitator superfamily Mef(E) transporter for macrolide resistance in Escherichia coli. Research in Microbiology 164, 226–235 (2013).

25. Tatsuno, I. et al. Functional Predominance of msr(D), Which Is More Effective as mef(A)-Associated Than mef(E)-Associated, Over mef(A)/mef(E) in Macrolide Resistance in Streptococcus pyogenes. Microbial Drug Resistance 24, 1089–1097 (2018).

26. Zhang, Y. et al. Predominant role of msr(D) over mef(A) in macrolide resistance in Streptococcus pyogenes. Microbiology (Reading) 162, 46–52 (2016).

27. Del Grosso, M., Camilli, R., Iannelli, F., Pozzi, G. & Pantosti, A. The mef(E)-Carrying Genetic Element (mega) of Streptococcus pneumoniae: Insertion Sites and Association with Other Genetic Elements. Antimicrob Agents Chemother 50, 3361–3366 (2006).

28. Del Grosso, M., Scotto d’Abusco, A., Iannelli, F., Pozzi, G. & Pantosti, A. Tn2009, a Tn916-Like Element Containing mef(E) in Streptococcus pneumoniae. Antimicrob Agents Chemother 48, 2037–2042 (2004).

29. Chancey, S. T. et al. Composite mobile genetic elements disseminating macrolide resistance in Streptococcus pneumoniae. Front. Microbiol. 6, (2015).

30. Iannelli, F. et al. Nucleotide sequence of conjugative prophage Φ1207.3 (formerly Tn1207.3) carrying the mef(A)/msr(D) genes for efflux resistance to macrolides in Streptococcus pyogenes. Front Microbiol 5, (2014).

31. González-Zorn, B. et al. Genetic basis for dissemination of armA. Journal of Antimicrobial Chemotherapy 56, 583–585 (2005).

32. Ambrose, K. D., Nisbet, R. & Stephens, D. S. Macrolide Efflux in Streptococcus pneumoniae Is Mediated by a Dual Efflux Pump (mel and mef) and Is Erythromycin Inducible. Antimicrob Agents Chemother 49, 4203–4209 (2005).

33. Chancey, S. T. et al. Transcriptional Attenuation Controls Macrolide Inducible Efflux and Resistance in Streptococcus pneumoniae and in Other Gram-Positive Bacteria Containing mef/mel(msr(D)) Elements. PLOS ONE 10, e0116254 (2015).

34. Ramu, H., Mankin, A. & Vazquez-Laslop, N. Programmed drug-dependent ribosome stalling. Mol Microbiol 71, 811–824 (2009).

35. Sothiselvam, S. et al. Macrolide antibiotics allosterically predispose the ribosome for translation arrest. Proc Natl Acad Sci U S A 111, 9804–9809 (2014).

36. de Block, T. et al. WGS of Commensal Neisseria Reveals Acquisition of a New Ribosomal Protection Protein (MsrD) as a Possible Explanation for High Level Azithromycin Resistance in Belgium. Pathogens 10, 384 (2021).

37. Lu, C. et al. Phenotypic and Genetic Characteristics of Macrolide and Lincosamide Resistant Ureaplasma urealyticum Isolated in Guangzhou, China. Curr Microbiol 61, 44–49 (2010).

38. Sharma, P. et al. Comparison of Antimicrobial Resistance and Pan-Genome of Clinical and Non-Clinical Enterococcus cecorum from Poultry Using whole-Genome Sequencing. Foods 9, 686 (2020).

39. Nikaido, H. Multidrug Resistance in Bacteria. Annual Review of Biochemistry 78, 119–146 (2009).

40. Nikaido, H. & Pagès, J.-M. Broad-specificity efflux pumps and their role in multidrug resistance of Gram-negative bacteria. FEMS Microbiol Rev 36, 340–363 (2012).

41. Datta, N., Hedges, R. w., Becker, D. & Davies, J. Plasmid-determined Fusidic Acid Resistance in the Enterobacteriaceae. Microbiology, 83, 191–196 (1974).

42. Kannan, K. et al. The general mode of translation inhibition by macrolide antibiotics. Proc Natl Acad Sci U S A 111, 15958–15963 (2014).

43. Vázquez-Laslop, N. & Mankin, A. S. How macrolide antibiotics work. Trends Biochem Sci 43, 668–684 (2018).

44. Murina, V. et al. ABCF ATPases Involved in Protein Synthesis, Ribosome Assembly and Antibiotic Resistance: Structural and Functional Diversification across the Tree of Life. Journal of Molecular Biology 431, 3568–3590 (2019).

45. Fürste, J. P. et al. Molecular cloning of the plasmid RP4 primase region in a multi-host-range tacP expression vector. Gene 48, 119–131 (1986).

46. Lutz, R. & Bujard, H. Independent and Tight Regulation of Transcriptional Units in Escherichia Coli Via the LacR/O, the TetR/O and AraC/I1-I2 Regulatory Elements. Nucleic Acids Res 25, 1203–1210 (1997).

47. Dubnau, D. Induction of ermC requires translation of the leader peptide. EMBO J 4, 533–537 (1985).

48. Dersch, P., Khan, M. A., Mühlen, S. & Görke, B. Roles of Regulatory RNAs for Antibiotic Resistance in Bacteria and Their Potential Value as Novel Drug Targets. Front. Microbiol. 8, (2017).

49. Banerjee, S., Chalissery, J., Bandey, I. & Sen, R. Rho-dependent Transcription Termination: More Questions than Answers. J Microbiol 44, 11–22 (2006).

50. Ray-Soni, A., Bellecourt, M. J. & Landick, R. Mechanisms of Bacterial Transcription Termination: All Good Things Must End. Annual Review of Biochemistry 85, 319–347 (2016).

51. Naville, M., Ghuillot-Gaudeffroy, A., Marchais, A. & Gautheret, D. ARNold: A web tool for the prediction of Rho-independent transcription terminators. RNA Biology 8, 11–13 (2011).

52. Takada, H. et al. Expression of Bacillus subtilis ABCF antibiotic resistance factor VmlR is regulated by RNA polymerase pausing, transcription attenuation, translation attenuation and (p)ppGpp. Nucleic Acids Res gkac497 (2022) doi:10.1093/nar/gkac497.

53. Yakhnin, A. V. et al. NusG controls transcription pausing and RNA polymerase translocation throughout the Bacillus subtilis genome. Proc Natl Acad Sci U S A 117, 21628–21636 (2020).

54. Shimizu, Y. et al. Cell-free translation reconstituted with purified components. Nat Biotechnol 19, 751–755 (2001).

55. Hartz, D., McPheeters, D. S., Traut, R. & Gold, L. Extension inhibition analysis of translation initiation complexes. in Methods in Enzymology vol. 164 419–425 (Academic Press, 1988).

56. Meydan, S. et al. Retapamulin-Assisted Ribosome Profiling Reveals the Alternative Bacterial Proteome. Molecular Cell 74, 481–493.e6 (2019).

57. Muto, H., Nakatogawa, H. & Ito, K. Genetically Encoded but Nonpolypeptide Prolyl-tRNA Functions in the A Site for SecM-Mediated Ribosomal Stall. Molecular Cell 22, 545–552 (2006).

58. Seip, B. & Innis, C. A. How Widespread is Metabolite Sensing by Ribosome-Arresting Nascent Peptides? Journal of Molecular Biology 428, 2217–2227 (2016).

59. Vázquez-Laslop, N. et al. Role of antibiotic ligand in nascent peptide-dependent ribosome stalling. Proc. Natl. Acad. Sci. U.S.A. 108, 10496–10501 (2011).

60. Hansen, J. L. et al. The Structures of Four Macrolide Antibiotics Bound to the Large Ribosomal Subunit. Molecular Cell 10, 117–128 (2002).

61. Mao, J. C. & Robishaw, E. E. Effects of macrolides on peptide-bond formation and translocation. Biochemistry 10, 2054–2061 (1971).

62. Poulsen, S. M., Kofoed, C. & Vester, B. Inhibition of the ribosomal peptidyl transferase reaction by the mycarose moiety of the antibiotics carbomycin, spiramycin and tylosin1 1Edited by D. E. Draper. Journal of Molecular Biology 304, 471–481 (2000).

63. Emmanuel, J. S., Sengupta, A., Gordon, E. R., Noble, J. T. & Cruz-Vera, L. R. The regulatory TnaC nascent peptide preferentially inhibits release factor 2-mediated hydrolysis of peptidyl-tRNA. J Biol Chem 294, 19224–19235 (2019).

64. Ohashi, Z. et al. Characterization of C+ located in the first position of the anticodon of Escherichia coli tRNAMet as N4-acetylcytidine. Biochimica et Biophysica Acta (BBA) - Nucleic Acids and Protein Synthesis 262, 209–213 (1972).

65. Varshney, U., Lee, C. P. & RajBhandary, U. L. From elongator tRNA to initiator tRNA. PNAS 90, 2305–2309 (1993).

66. Pierson, W. E. et al. Uniformity of Peptide Release Is Maintained by Methylation of Release Factors. Cell Reports 17, 11–18 (2016).

67. Schmeing, T. M., Huang, K. S., Strobel, S. A. & Steitz, T. A. An induced-fit mechanism to promote peptide bond formation and exclude hydrolysis of peptidyl-tRNA. Nature 438, 520–524 (2005).

68. Arenz, S. et al. Molecular basis for erythromycin-dependent ribosome stalling during translation of the ErmBL leader peptide. Nature Communications 5, 3501 (2014).

69. Lovmar, M. et al. The Molecular Mechanism of Peptide-mediated Erythromycin Resistance *. Journal of Biological Chemistry 281, 6742–6750 (2006).

70. Tenson, T., Xiong, L., Kloss, P. & Mankin, A. S. Erythromycin Resistance Peptides Selected from Random Peptide Libraries*. Journal of Biological Chemistry 272, 17425–17430 (1997).

71. Koch, M., Willi, J., Pradère, U., Hall, J. & Polacek, N. Critical 23S rRNA interactions for macrolide-dependent ribosome stalling on the ErmCL nascent peptide chain. Nucleic Acids Res 45, 6717–6728 (2017).

72. Vázquez-Laslop, N., Ramu, H., Klepacki, D., Kannan, K. & Mankin, A. S. The key function of a conserved and modified rRNA residue in the ribosomal response to the nascent peptide. EMBO J 29, 3108–3117 (2010).

73. Vimberg, V. et al. Ribosome-Mediated Attenuation of vga(A) Expression Is Shaped by the Antibiotic Resistance Specificity of Vga(A) Protein Variants. Antimicrobial Agents and Chemotherapy 64, (2020).

74. Jacquet, E. et al. ATP Hydrolysis and Pristinamycin IIA Inhibition of the Staphylococcus aureus Vga(A), a Dual ABC Protein Involved in Streptogramin A Resistance *. Journal of Biological Chemistry 283, 25332–25339 (2008).

75. Mayford, M. & Weisblum, B. Conformational alterations in the ermC transcript in vivo during induction. EMBO J 8, 4307–4314 (1989).

76. Dar, D. et al. Term-seq reveals abundant ribo-regulation of antibiotics resistance in bacteria. Science 352, aad9822 (2016).

77. Koberska, M. et al. Beyond Self-Resistance: ABCF ATPase LmrC Is a Signal-Transducing Component of an Antibiotic-Driven Signaling Cascade Accelerating the Onset of Lincomycin Biosynthesis. mBio 12, e0173121 (2021).

78. Ohki, R., Tateno, K., Takizawa, T., Aiso, T. & Murata, M. Transcriptional Termination Control of a Novel ABC Transporter Gene Involved in Antibiotic Resistance in Bacillus subtilis. J Bacteriol 187, 5946–5954 (2005).

79. Dunkle, J. A., Xiong, L., Mankin, A. S. & Cate, J. H. D. Structures of the Escherichia coli ribosome with antibiotics bound near the peptidyl transferase center explain spectra of drug action. Proc Natl Acad Sci U S A 107, 17152–17157 (2010).

80. Desai, N. et al. Elongational stalling activates mitoribosome-associated quality control. Science 370, 1105–1110 (2020).

81. Golubev, A. et al. Cryo-EM structure of the ribosome functional complex of the human pathogen Staphylococcus aureus at 3.2 Å resolution. FEBS Letters 594, 3551–3567 (2020).

82. Li, W. et al. Structural basis for selective stalling of human ribosome nascent chain complexes by a drug-like molecule. Nat Struct Mol Biol 26, 501–509 (2019).

83. Croucher, N. J. et al. Rapid pneumococcal evolution in response to clinical interventions. Science 331, 430–434 (2011).

84. Miller, J. R. GraphPad Prism Version 4.0 Step-by-Step Examples. vol. GraphPad Software Inc., San Diego (GraphPad Software Inc., San Diego, 2003).

85. Stead, M. B. et al. RNAsnap™: a rapid, quantitative and inexpensive, method for isolating total RNA from bacteria. Nucleic Acids Res 40, e156 (2012).

86. Seefeldt, A. C. et al. The proline-rich antimicrobial peptide Onc112 inhibits translation by blocking and destabilizing the initiation complex. Nature Structural & Molecular Biology 22, 470–475 (2015).

87. Zheng, S. Q. et al. MotionCor2 - anisotropic correction of beam-induced motion for improved cryo-electron microscopy. Nat Methods 14, 331–332 (2017).

88. Zhang, K. Gctf: Real-time CTF determination and correction. J Struct Biol 193, 1–12 (2016).

89. Zivanov, J. et al. New tools for automated high-resolution cryo-EM structure determination in RELION-3. eLife 7, e42166 (2018).

90. Kucukelbir, A., Sigworth, F. J. & Tagare, H. D. The Local Resolution of Cryo-EM Density Maps. Nat Methods 11, 63–65 (2014).

91. Emsley, P. & Cowtan, K. Coot: model-building tools for molecular graphics. Acta Cryst D 60, 2126–2132 (2004).

92. Liebschner, D. et al. Macromolecular structure determination using X-rays, neutrons and electrons: recent developments in Phenix. Acta Cryst D 75, 861–877 (2019).

93. Boccaletto, P. et al. MODOMICS: a database of RNA modification pathways. 2017 update. Nucleic Acids Res 46, D303–D307 (2018).

94. Bourgeois, G. et al. Structural basis of the interaction between cyclodipeptide synthases and aminoacylated tRNA substrates. RNA 26, 1589–1602 (2020).

95. Waterhouse, A. M., Procter, J. B., Martin, D. M. A., Clamp, M. & Barton, G. J. Jalview Version 2—a multiple sequence alignment editor and analysis workbench. Bioinformatics 25, 1189–1191 (2009).

96. Schindelin, J. et al. Fiji: an open-source platform for biological-image analysis. Nature Methods 9, 676–682 (2012).

97. Goddard, T. D. et al. UCSF ChimeraX: Meeting modern challenges in visualization and analysis. Protein Sci 27, 14–25 (2018).

98. Pettersen, E. F. et al. UCSF Chimera—A visualization system for exploratory research and analysis. Journal of Computational Chemistry 25, 1605–1612 (2004).

