## Supplementary material for "Regulation of the macrolide resistance ABC-F translation factor MsrD": SI_Fostier_et_al_Aug_2022

**Supplementary Table 1. Minimum inhibitory concentration (MIC) and half maximal inhibitory concentration ($IC_{50}$) of *E. coli* DB10 expressing *msrD* variants in presence of erythromycin.** See Methods for experimental details. MIC values exceeding control plasmid are shown in bold.

| | **Erythromycin** | |
|---|---|---|
| ***E. coli* DB10** | **MIC (µM)** | **$IC_{50}$ (µM)** |
| pBAD-*Control* | 2 | 0,179 ± 0,007 |
| pBAD-*msrD*$_{WT}$ | **16** | 4,592 ± 0,582 |
| pBAD-*msrD*$_{EQ2}$ | 2 | 0,267 ± 0,027 |
| pBAD-*msrD*$_{E125Q}$ | **4** | 1,427 ± 0,147 |
| pBAD-*msrD*$_{E434Q}$ | 2 | 0,342 ± 0,04 |
| pBAD-*msrD*$_{\Delta Loop}$ | 2 | 0,209 ± 0,025 |
| pBAD-*msrD*$_{\Delta PtIM}$ | 2 | 0,295 ± 0,048 |
| pBAD-*msrD*$_{R241A}$ | **16** | 3,632 ± 0,316 |
| pBAD-*msrD*$_{L242A}$ | **8** | 1,621 ± 0,152 |
| pBAD-*msrD*$_{H244A}$ | **8** | 1,492 ± 0,126 |
| pBAD-*msrD*$_{H244W}$ | **4** | 0,446 ± 0,057 |

**Supplementary Table 2. Cryo-EM data collection and refinement statistics.**

| | **Erythromycin-stalled *Escherichia coli* 70S ribosome with streptococcal MsrDL nascent chain** (PDB 7Q4K, EMD-13805, EMD-13806, EMD-13807, EMD-13808) |
|---|---|
| **Data collection and processing** | |
| Microscope | FEI Talos Arctica (IECB, Pessac, France) |
| Detector | K2 Summit direct electron detector (Gatan) |
| Magnification (X) | 120,000 |
| Voltage (kV) | 200 |
| Electron exposure ($e^{-}$/Å$^2$) | 64 |
| Defocus range (µm) | -0.5 to -2.7 |
| Pixel size (Å) | 1.2 |
| Symmetry imposed | C1 |
| Initial particle images (no.) | 158,200 |
| Final particle images (no.) | 62,093 |
| Map resolution (Å) | 70S ribosome (EMD-13805): 3<br>50S subunit (EMD-13806): 2.97<br>30S subunit Body (EMD-13807): 3.08<br>30S subunit Head (EMD-13808): 3.3 |
| FSC threshold | 0.143 |
| **Model building and refinement** | |
| Initial model (PDB code) | 6TC3 |
| Model resolution (Å) | 2.7 |
| FSC threshold | 0.143 |
| Model resolution range (Å) | 2.5 – 8.7 |
| Model composition | - |
| Non-hydrogen atoms | 146,618 |
| Protein residues | 5,682 |
| RNA bases | 4,710 |

| | |
|---|---|
| Ligands | 173 |
| B-factor (Å$^2$) | - |
| Protein (min./max./mean) | 31.70/183.09/69.05 |
| Ligands (min./max./mean) | 20.00/458.82/75.79 |
| R.m.s. deviations | - |
| Bond lengths (Å) | 0.015 |
| Bond angles (°) | 1.648 |
| Validation | - |
| MolProbity score | 1.49 |
| Clashscore | 3.83 |
| Poor rotamers (%) | 0.63 |
| Ramachandran plot | - |
| Favored (%) | 95.42 |
| Allowed (%) | 4.45 |
| Disallowed (%) | 0.13 |

**Supplementary Table 3. Strains and plasmids used in this study.**

| **Bacterial strains** | **References** |
|---|---|
| *E. coli* DB10 (Derived from *E. coli* PR7) | [1] |
| *E. coli* K12 MG1655 | N/A |
| **Plasmids** | **References** |
| pBAD-*Control* (pBAD33) | [2] |
| pBAD-*msrD*$_{\mathrm{WT}}$ (pVN50) | [2] |
| pBAD-*msrD*$_{\mathrm{EQ2}}$ | This study |
| pBAD-*msrD*$_{\mathrm{E125Q}}$ | This study |
| pBAD-*msrD*$_{\mathrm{E434Q}}$ | This study |
| pBAD-*msrD*$_{\Delta \mathrm{Loop}}$ | This study |
| pBAD-*msrD*$_{\Delta \mathrm{PtIM}}$ | This study |
| pBAD-*msrD*$_{\mathrm{R241A}}$ | This study |
| pBAD-*msrD*$_{\mathrm{L242A}}$ | This study |
| pBAD-*msrD*$_{\mathrm{H244A}}$ | This study |
| pBAD-*msrD*$_{\mathrm{H244W}}$ | This study |
| pMMB-67EH | [3] |
| pMMB-67EH-*yfp* | This study |
| pMMBpLlacO-1-67EH-*yfp* | This study |
| pMMB-*msrDL*-*msrD*$_{(1-3)}$*:yfp* | This study |
| pMMB-*msrDL*$_{(\mathrm{no\_ORF})}$-*msrD*$_{(1-3)}$*:yfp* | This study |
| pMMB-*msrDL*$_{(\mathrm{no\_term})}$-*msrD*$_{(1-3)}$*:yfp* | This study |
| pMMB-*msrDL*$_{(\mathrm{Y2A})}$-*msrD*$_{(1-3)}$*:yfp* | This study |
| pMMB-*msrDL*$_{(\mathrm{L3A})}$-*msrD*$_{(1-3)}$*:yfp* | This study |
| pMMB-*msrDL*$_{(\mathrm{I4A})}$-*msrD*$_{(1-3)}$*:yfp* | This study |
| pMMB-*msrDL*$_{(\mathrm{F5A})}$-*msrD*$_{(1-3)}$*:yfp* | This study |
| pMMB-*msrDL*$_{(\mathrm{M6A})}$-*msrD*$_{(1-3)}$*:yfp* | This study |
| pMMB-*msrDL*$_{(\mathrm{UAA>UGA})}$-*msrD*$_{(1-3)}$*:yfp* | This study |
| pMMB-*msrDL*$_{(\mathrm{UAA>UAG})}$-*msrD*$_{(1-3)}$*:yfp* | This study |
| pMMB-*msrDL*$_{(\mathrm{WT\text{-}isocodons})}$-*msrD*$_{(1-3)}$*:yfp* | This study |
| pMMB-*msrDL*$_{(\mathrm{MYLIFMA\text{-}isocodons})}$-*msrD*$_{(1-3)}$*:yfp* | This study |

**Supplementary Table 4. Oligonucleotides used in this study.**

| No. | Name | Sequence (5' → 3') | Purpose |
|---|---|---|---|
| **pBAD plasmids** | | | |
| 1 | msrD_F | ATGGAATTAATATTAAAAGCAAAAGACATTCGTGTGG | Amplification of *msrD*$_{WT}$ |
| 2 | msrD_R | TTAGTGATGGTGATGGTGATGTTTCAGATTTATTTTCTTATC | |
| 3 | pBAD_F | CATCACCATCACCATCACTAATCTAGAGTCGACCTGCAGGC | Amplification of pBAD backbone |
| 4 | pBAD_R | GCTTTTAATATTAATTCCATGGTGAATTCCTCCTGCTAGCC | |
| 5 | msrD$_{EQ2}$_F1 | GGTATTTTAGCGGATCAACCTACGAG | *msrD* mutagenesis |
| 6 | msrD$_{EQ2}$_R1 | GGAAGTTACTGGGTTGATCCATTATTAG | |
| 7 | msrD$_{EQ2}$_F2 | AACCCAGTAACTTCCTTGACATACC | |
| 8 | msrD$_{EQ2}$_R2 | GATCCGCTAAAATACCATGAACC | |
| 9 | msrD$_{E125Q}$_F | GGTATTTTAGCGGATCAACCTACGAGCCATTTAG | |
| 10 | msrD$_{E125Q}$_R | GGCTCGTAGGTTGATCCGCTAAAATACCATG | |
| 11 | msrD$_{E434Q}$_F | CTAATAATGGATCAACCCAGTAACTTCCTTGAC | |
| 12 | msrD$_{E434Q}$_R | GGAAGTTACTGGGTTGATCCATTATTAGGATG | |
| 13 | msrD$_{\Delta Loop}$_F | GGAAAGGGCTGCGGAGGAAAAGGGAGGAGGAAAGATGTATAATGCTGCTAAAAC | |
| 14 | msrD$_{\Delta Loop}$_R | GCAGCATTATACATCTTTCCTCCTCCCTTTTCCTCCGCAGCCCTTTCCAATCGGG | |
| 15 | msrD$_{\Delta PtIM}$_F | CTGATTATCTTCGTCAGAAAGGAGGAGGACCGGAAGGCATTCGCAGAATTCG | |
| 16 | msrD$_{\Delta PtIM}$_R | CGAATTCTGCGAATGCCTTCCGGTCCTCCTCCTTTCTGACGAAGATAATCAGAATAG | |
| 17 | msrD$_{R241A}$_F | GAAGACGGAGGGGCTTTAGCTCATCAAAAATC | |
| 18 | msrD$_{R241A}$_R | GATGAGCTAAAGCCCCTCCGTCTTCAGTAC | |
| 19 | msrD$_{L242A}$_F | GACGGAGGGCGTGCAGCTCATCAAAAATCAATAG | |
| 20 | msrD$_{L242A}$_R | GATTTTTGATGAGCTGCACGCCCTCCGTCTTCAG | |
| 21 | msrD$_{H244A}$_F | GCGTTTAGCTGCTCAAAAATCAATAGGAAGTAAGG | |

| | | | |
|---|---|---|---|
| 22 | $msrD_{H244A}$_R | CTATTGATTTTTGAGCAGCTAA ACGCCCTCCGTCTTC | |
| 23 | $msrD_{H244W}$_F | GCGTTTAGCTTGGCAAAAATCA ATAGGAAGTAAGG | |
| 24 | $msrD_{H244W}$_R | CTATTGATTTTTGCCAAGCTAA ACGCCCTCCGTCTTC | |
| **pMMB plasmids** | | | |
| 25 | pMMB-PlacO_F | GTATAAATGTGAGCGGATAAC ATTGACATTGTGAGCGGATAA CAAGATACTGAGCACATCACA CAGGAAACAGAATATGTCC | Replacement of $P_{tac}$ promoter by $P_{lacO}$ [4] |
| 26 | pMMB-PlacO_R | GTTATCCGCTCACATTTATACA GCTCATTTCAGAATATTTGCC | |
| 27 | msrDL-$msrD_{(1-3)}$:yfp_F1 | CGCAGGGTTTTCCCTGCATAC AAGCAAATGAAAGCATGCGAT TATAGACAGGAGGAAATGTTAT GGAATTAATCGTAAAAATCGTG AGCAAGG | Fusion of *msrDL-msrD*$_{(1-3)}$ cistron to *yfp* |
| 28 | msrDL-$msrD_{(1-3)}$:yfp_F2 | CTGAGCACAACAATATTGGAG GAATATTTATGTATCTTATTTTC ATGTAACTCTTCCTGCTAAAAT CGCAGGGTTTTCCCTGCATAC AAGC | |
| 29 | yfp_R | TTACTTGTACAGCTCGTCCATG CCGAGAGTGATCCCGGCGGC GG | |
| 30 | pMMB-backbone_F | CATGGACGAGCTGTACAAGTA ATAATTCGAGCTCGGTACCCG GG | - |
| 31 | pMMB-backbone_R | CCTCCAATATTGTTGTGCTCAG TATCTTGTTATCCGCTCACAAT GTC | - |
| 32 | pMMB-control_F | CAACAATATTGGAGGAATATTT TAATTCGAGCTCGGTACCC | - |
| 33 | pMMB-control_R | GGGTACCGAGCTCGAATTAAA ATATTCCTCCAATATTGTTG | - |
| 34 | $msrDL_{(no_term)}$_F | ATGTATCTTATTTTCATGTAACT CTCCCTGCATACAAGCAAATG | Deletion of RIT |
| 35 | $msrDL_{(no_term)}$_R | AGAGTTACATGAAAATAAGATA CATAAATATTCCTCC | |
| 36 | $msrDL_{(no_ORF)}$_F | CAACAATATTGGAGGAATATTT TAGTATCTTATTTTCATGTAACT CTTCC | Suppression of ORF *msrDL* |
| 37 | $msrDL_{(no_ORF)}$_R | GGAAGAGTTACATGAAAATAA GATACTAAAATATTCCTCCAAT ATTGTTG | |
| 38 | $msrDL_{Y2A}$_F | CAATATTGGAGGAATATTTATG GCACTTATTTTCATGTAACTCT TCC | *msrDL* mutagenesis |
| 39 | $msrDL_{Y2A}$_R | GGAAGAGTTACATGAAAATAA GTGCCATAAATATTCCTCCAAT ATTG | |
| 40 | $msrDL_{L3A}$_F | TTGGAGGAATATTTATGTATGC AATTTTCATGTAACTCTTCC | |

| | | | |
|---|---|---|---|
| 41 | msrDL$_{L3A}$_R | GGAAGAGTTACATGAAAATTGCATACATAAATATTCCTCCAA | |
| 42 | msrDL$_{I4A}$_F | GGAGGAATATTTATGTATCTTGCATTCATGTAACTCTTCCTG | |
| 43 | msrDL$_{I4A}$_R | CAGGAAGAGTTACATGAATGCAAGATACATAAATATTCCTCC | |
| 44 | msrDL$_{F5A}$_F | GGAATATTTATGTATCTTATTGCAATGTAACTCTTCCTG | |
| 45 | msrDL$_{F5A}$_R | CAGGAAGAGTTACATTGCAATAAGATACATAAATATTCC | |
| 46 | msrDL$_{M6A}$_F | ATTTATGTATCTTATTTTCGCATAACTCTTCCTGCTAAAATCGCAGG | |
| 47 | msrDL$_{M6A}$_R | CCTGCGATTTTAGCAGGAAGAGTTATGCGAAAATAAGATACATAAAT | |
| 48 | msrDL$_{TAG}$_F | GTATCTTATTTTCATGTAGCTCTTCCTGCTAAAATCG | |
| 49 | msrDL$_{TAG}$_R | CGATTTTAGCAGGAAGAGCTACATGAAAATAAGATAC | |
| 50 | msrDL$_{TGA}$_F | GTATCTTATTTTCATGTGACTCTTCCTGCTAAAATCG | |
| 51 | msrDL$_{TGA}$_R | CGATTTTAGCAGGAAGAGTCACATGAAAATAAGATAC | |
| 52 | msrDL$_{MYLIFMA}$-isocodons_F | ATGTACCTGATCTTCATGGCCTAACTCTTCCTGCTAAAATCGCAGGG | |
| 53 | msrDL$_{MYLIFMA}$-isocodons_R | TTAGGCCATGAAGATCAGGTACATAAATATTCCTCCAATATTGTTGTGCTCAGTATCTTGTTATCCGC | |
| 54 | msrDL$_{WT}$-isocodons_F | ATTTATGTACCTGATCTTCATGTAACTCTTCCTGCTAAAATCGC | |
| 55 | msrDL$_{WT}$-isocodons_R | GCGATTTTAGCAGGAAGAGTTACATGAAGATCAGGTACATAAAT | |
| **In vitro transcription and translation** | | | |
| 56 | T7_pMMB_F | GCGAATTAATACGACTCACTATAGGGAGCGGATAACAAGATACTGAGCAC | - |
| 57 | TP_msrDL_R | GGTTATAATGAATTTTGCTTATTTAATTCCATAACATTTCCTCC | - |
| 58 | TP_NV1_R | GGTTATAATGAATTTTGCTTATT | CY5 chromophore modification in 5’ |
| **Northern blot probe** | | | |
| 59 | pMMB_3UTR | CAGCCAAGCTTGCATGCCTGCAGGTCGACTCTAGAGGATCCCCGGG | - |
| **Cryo-EM** | | | |
| 60 | cryoEM-msrDL_R | GCAGGGAAAACCCTGCG | - |

**Supplementary Table 5. DNA templates used in this study.**

| No. | Name | Sequence (5' → 3') |
|---|---|---|
| 1 | TP_msrDL$_{WT}$ | GCGAATTAATACGACTCACTATAGGGAGCGGATAACAAGATACTGAGCACAACAATATTGGAGGAATATTTATGTATCTTATTTTCATGTAACTCTTCCTGCTAAAATCGCAGGGTTTTCCCTGCATACAAGCAAATGAAAGCATGCGATTATAGACAGGAGGAAATGTTATGGAATTAAATAAGCAAAATTCATTATAACC |
| 2 | TP_msrDL$_{7A\text{-}iso}$ | GCGAATTAATACGACTCACTATAGGGAGCGGATAACAAGATACTGAGCACAACAATATTGGAGGAATATTTATGTACCTGATCTTCATGGCCTAACTCTTCCTGCTAAAATCGCAGGGTTTTCCCTGCATACAAGCAAATGAAAGCATGCGATTATAGACAGGAGGAAATGTTATGGAATTAAATAAGCAAAATTCATTATAACC |
| 3 | CryoEM_*msrDL* | GCGAATTAATACGACTCACTATAGGGAGCGGATAACAAGATACTGAGCACAACAATATTGGAGGAATATTTATGTATCTTATTTTCATGTAACTCTTCCTGCTAAAATCGCAGGGTTTTCCCTGC |

### SUPPLEMENTARY FIGURE 1

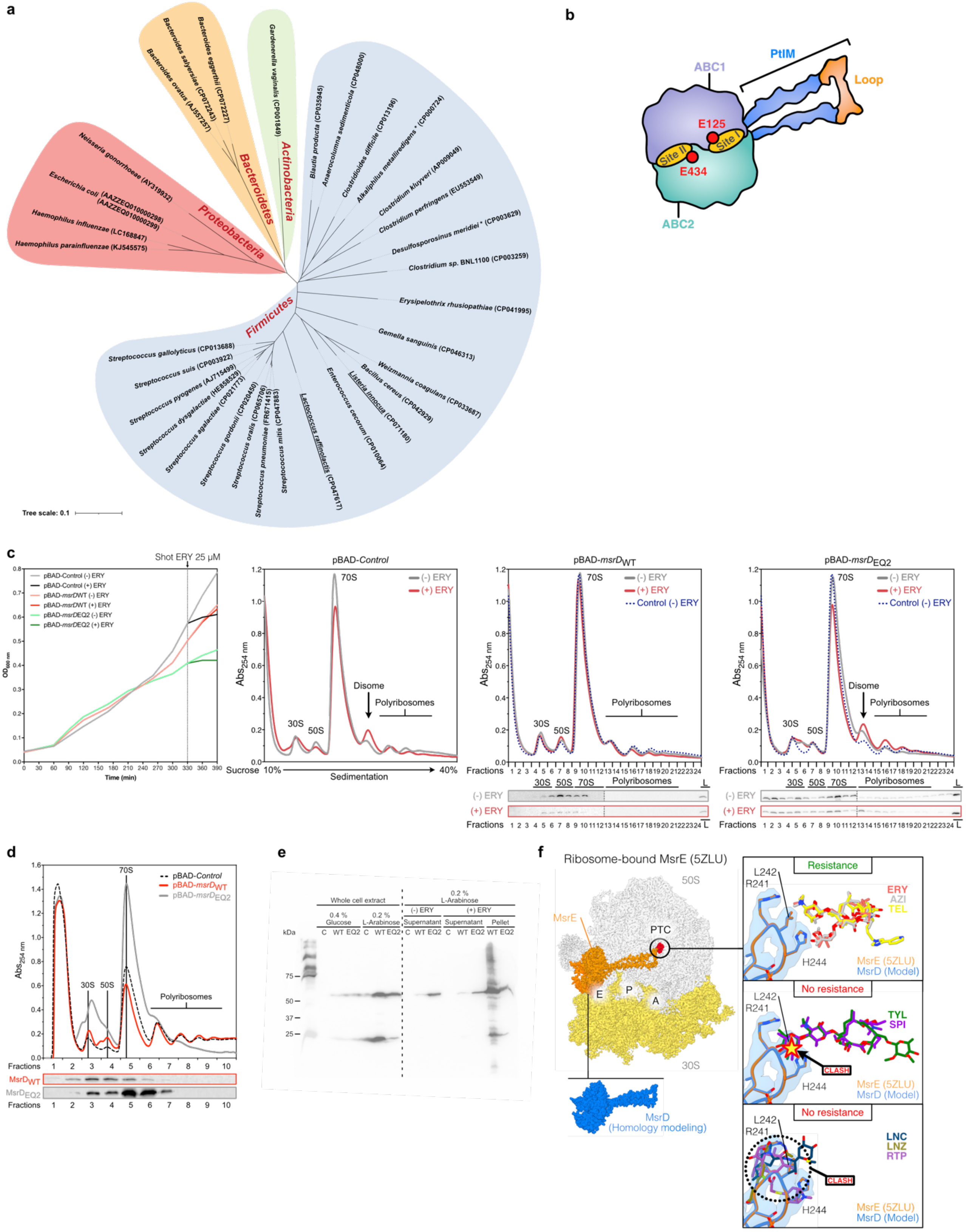

**Supplementary figure 1. *In vivo* characterization of MsrD**. (a) Dissemination of the *mefA/msrD* macrolide resistance operon among a wide range of non-pathogenic and pathogenic bacterial species. The *mefA/msrD* operon (Genbank accession No. FR671415) was blasted and identified in indicated species, integrated to the genome. Species where the operon was found on plasmid are underlined. Species where *msrD* was found to disseminate alone with *msrDL* are indicated by an asterisk. Corresponding Genebank accession numbers are indicated between brackets. To generate the tree, 16S rRNA sequences were retrieved, aligned using Clustal W [5], and resulting cladogram was adapted on iTOL (https://itol.embl.de/). Colored zones indicate bacterial phyla. (b) Schematic of MsrD illustrating main features of ABC-F translation factors and the position of ATP hydrolysis Site I and Site II whose catalytic residues are respectively E125 and E434. Colors are the same as Fig. 1b. (c) Growth curves, polyribosomes analysis and western blotting of *E. coli* DB10 strain expressing *msrD* (pBAD-$msrD_{WT}$), $msrD_{EQ2}$ (pBAD-$msrD_{EQ2}$) or a control (pBAD-*Control*) unexposed or exposed to 25 µM of ERY during one hour (Cells were treated after 330 min as indicated by “Shot ERY 25 µM”). “L” stands for total lysate. Note that these sample were precipitated with trichloroacetic acid (d) Polyribosomes analysis and western-blotting of *E. coli* K12 MG1655 expressing *msrD*WT (pBAD-*msrD*WT), *msrD*EQ2 (pBAD-*msrD*EQ2) or a control (pBAD-*Control*). As no erythromycin was added, 100 µg.ml$^{-1}$ chloramphenicol was added to stabilize translating ribosomes. (e) Solubility assay shows that $MsrD_{WT}$ tends precipitate in insoluble fraction while $MsrD_{EQ2}$ tends to stay more soluble. *E. coli* DB10 strain containing pBAD-*Control* (“C”), pBAD-$msrD_{WT}$ (WT) or pBAD-$msrD_{EQ2}$ (“EQ2”) was inoculated at $OD_{600}$=0.1 and grew in presence of 0.4 % Glucose or 0.2 % L-Arabinose at 37 °C under vigorous shaking for 6 hours. Cultures were normalized, centrifuged and resuspended (1 ml at $OD_{600}$=1 was resuspended in 30 µL Laemmli 1X). In total, 5 µL of whole cell extract were loaded on SDS-PAGE gel. In parallel, cultures similar to those realized for polyribosome analysis presented in Fig. 1c and 1d were done. Briefly, cells were inoculated at $OD_{600}$=0.1 and grew in presence of 0.2 % L-Arabinose at 37 °C under vigorous shaking. During mid-exponential phase, 25 µM ERY was added or not. Cells were harvested and lysed after 1 h, lysate being clarified by centrifugation from insoluble fraction. After normalization of lysate, a total of 10 µg of RNAs were load on SDS-PAGE gel. Western-blotting was performed as explained in Methods. (f) Steric occlusion mechanism by MsrD is incompatible with its resistance profile. A homology model of MsrD was generated using SWISS-MODEL [6] and aligned to ribosome-bound MsrE structure (PDB 5ZLU) [7]. Antibiotics to which MsrD provides resistance (ERY PDB: 6ND6, red; AZI PDB: 4V7Y, gray; TEL PDB: 4V7Z, yellow) or not (TYL PDB: 1K9M, green; SPI PDB: 1KD1, violet; LNC PDB: 5HKV, dark teal; LNZ PDB: 3CPW, khaki; RTP PDB: 2OGO, purple) were aligned based on domain V of 23S rRNA [8–13]. Density for MsrE is shown in pale blue, and conserved residues R241A, L242A and H244A are indicated.

#### SUPPLEMENTARY FIGURE 2

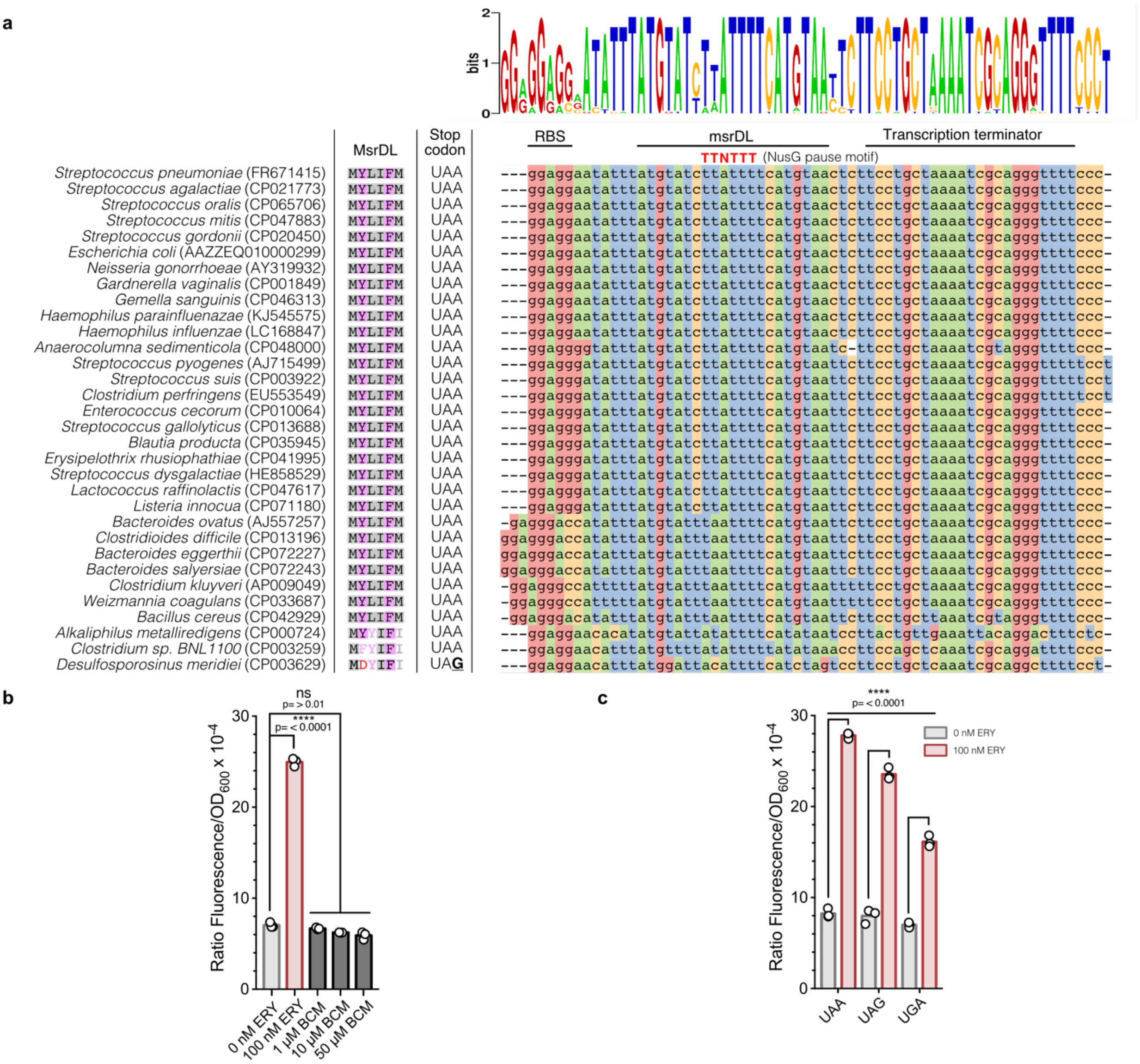


**Supplementary figure 2. Rho-independent transcription termination regulates *msrD* expression**. (a) Conservation of *msrDL*, NusG-dependent RNAP pausing site and its rho-independent transcription terminator. Sequences corresponding to Fig. 1a were aligned with Clustal W [5], and visualized with JalView according to nucleotide [14]. Genbank accession numbers are indicated between brackets. Logo was generated using WebLogo (https://weblogo.berkeley.edu/logo.cgi). (b) Bicyclomycin (BCM) failed to constitutively induce *msrD*$_{(1\text{-}3)}$:*yfp* expression in absence of ERY after 17 h, demonstrating that *msrD* is not regulated by a Rho-dependent terminator. Error bars represent mean ± s.d. for triplicate experiments. (c) Effects of *msrDL* stop codon mutation on the expression of *msrD*$_{(1:3):}$*yfp*. Bacteria were grown during 17 h in presence of 1 mM IPTG, in the absence (grey histograms) or in the presence of 100 nM ERY (red histograms). Error bars represent mean ± s.d. for triplicate experiments.

#### SUPPLEMENTARY FIGURE 3

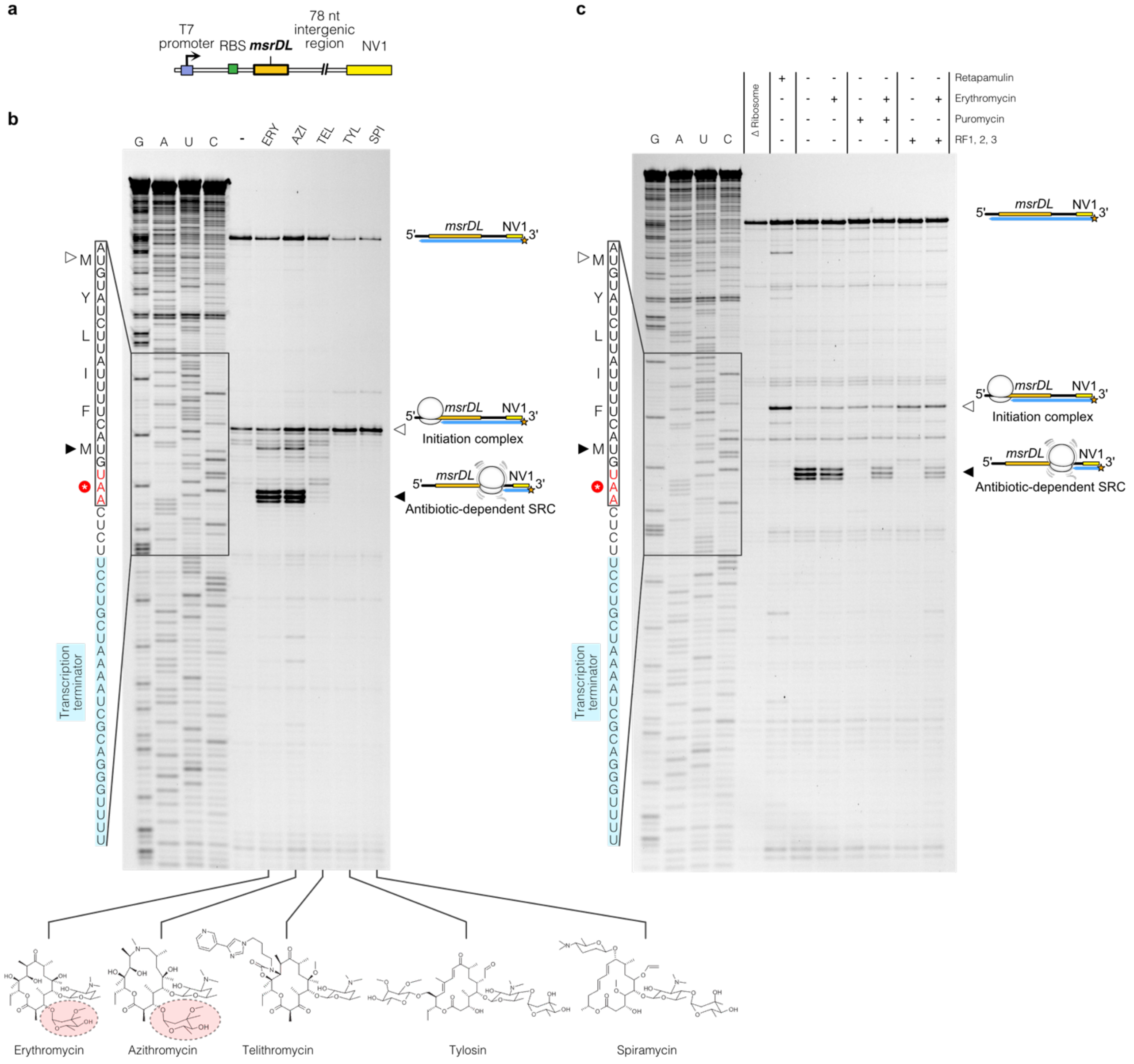


**Supplementary figure 3. Biochemical characterization of MsrDL mode of action**. (a) Schematic of the matrix TP_msrDL$_{WT}$ (Supplementary Table 5) used to generate synthetic mRNAs for *in vitro* experiments (RBS, ribosome binding site; NV1, annealing site for CY5-labelled primer). (b and c) Uncropped toe-printing gels presented in Fig. 3a and 3c. Formation of MsrDL-SRC would occlude formation of the Rho-independent transcription terminator (shown in light blue).

#### SUPPLEMENTARY FIGURE 4

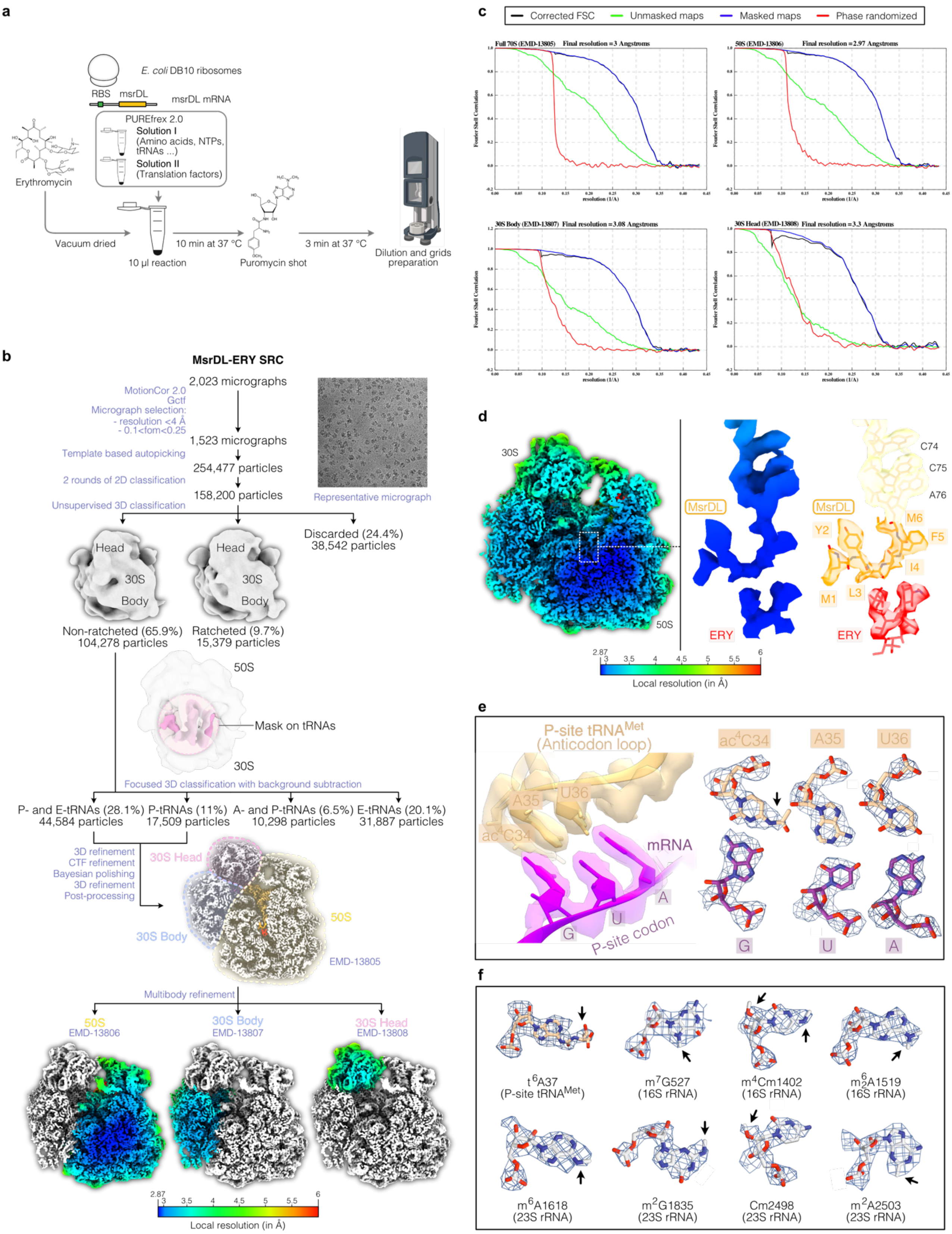

**Supplementary figure 4. Generation of the cryo-EM sample, data processing and model building**. (a) Workflow for the generation of cryo-EM sample. See Methods for details. (b) Cryo-EM data processing workflow as described in the Methods. EMD accession numbers are indicated for each map. Maps obtained after multibody refinement are shown as transverse sections and colored according to local resolution. (c) Fourier Shell Correlation (FSC) curves of the 70S ribosome and the maps obtained after multibody refinement. (d) Transverse section of the composite map obtained after multibody refinement and isolated densities for MsrDL-tRNA and ERY, colored according to local resolution. The corresponding atomic reconstruction of the nascent chain is shown beside. (e) Details of the mRNA codon-tRNA anticodon interactions and corresponding densities for each nucleotide. (f) Details of clearly identifiable post-transcriptional modifications and corresponding densities.

#### SUPPLEMENTARY FIGURE 5

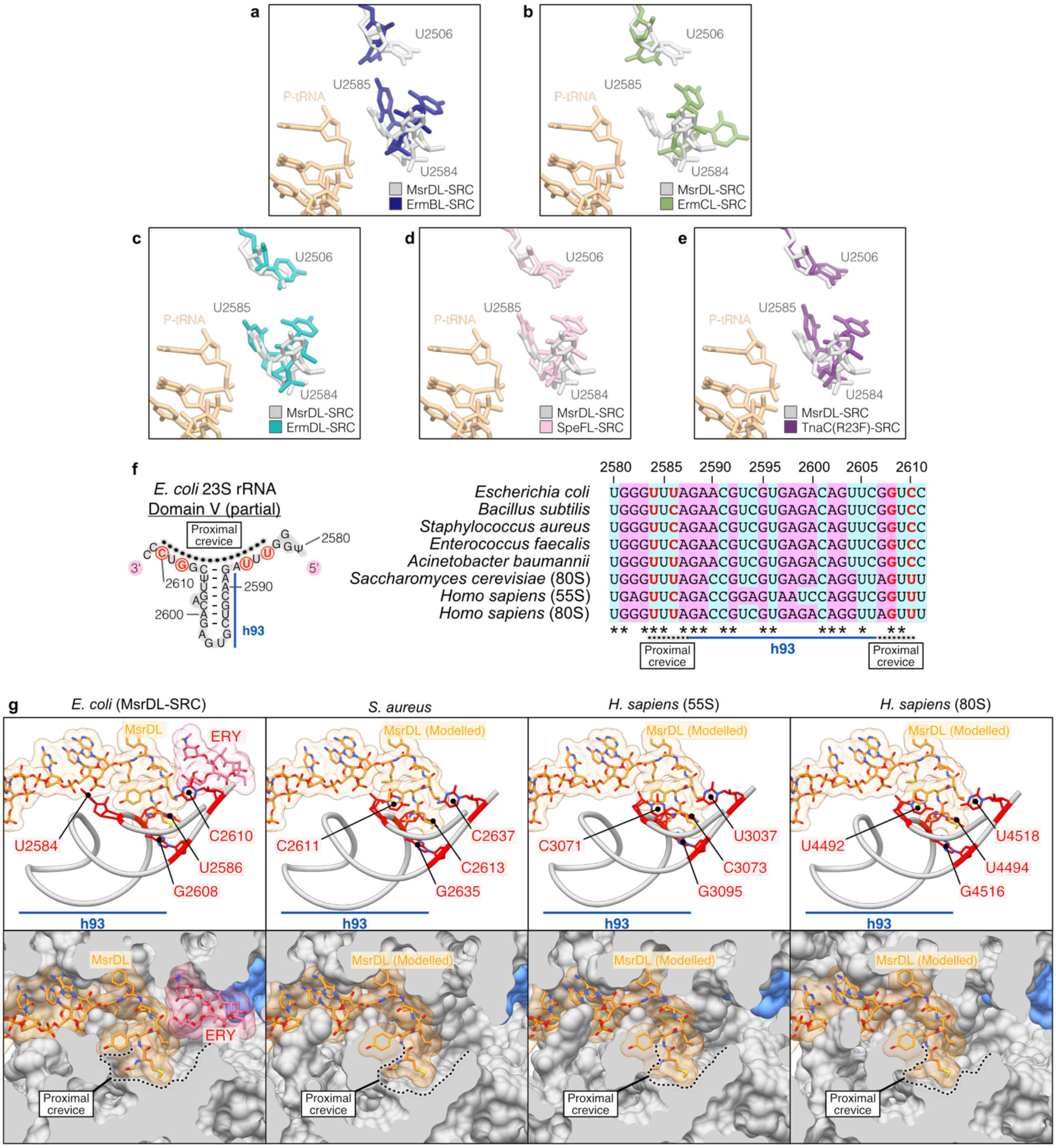

**Supplementary figure 5. MsrDL engages within a universally conserved crevice at the NPET entrance.** (a to e) Comparison of the conformation of 23S rRNA bases U2506, U2584 and U2585 for MsrDL-, ErmBL-, ErmCL-, ErmDL-, SpeFL- and TnaC(R23F)-stalled ribosomes structures [15–19](respectively PDB: 5JTE, 3J7Z, 7NSO, 6TC3, 7O1A). Structures were aligned based on domain V of 23S rRNA. (f) Sequence conservation of the proximal crevice between far-related species. The diagram shows a part of 23S rRNA domain V of *E. coli* and the location of proximal crevice at the base of h93. Large subunit rRNAs were aligned using Clustal W [5]

and visualized with Jalview [14] (purines, purple; pyrimidines, teal). Bases delimitating the proximal crevice (U2584, U2586, G2608 and C2610) are highlighted in red. Nucleotides numbering is relative to *E. coli* 23S rRNA sequence. (g) Structural conservation of the proximal crevice between far-related species. Top, cartoon representation of h93 and proximal crevice in *E. coli*, *S. aureus*, *H. sapiens* 55S mitoribosome and 80S cytosolic ribosome [20–22] (respectively PDB: 6YEF, 7A5F, 6OLI). Bases delimitating the proximal crevice are highlighted in red and numbered relative to the considered specie. Bottom, sagittal cut of ribosome tunnel shown on top panels depicted as surface. Ribosomal protein uL4 is shown in pale blue. Structures were aligned based on domain V of 23S rRNA.

**SUPPLEMENTARY REFERENCES**


1. Datta, N., Hedges, R. W., Becker, D. & Davies, J. Plasmid-determined Fusidic Acid Resistance in the Enterobacteriaceae. *Microbiology,* **83**, 191–196 (1974).
2. Nunez-Samudio, V. & Chesneau, O. Functional interplay between the ATP binding cassette Msr(D) protein and the membrane facilitator superfamily Mef(E) transporter for macrolide resistance in Escherichia coli. *Research in Microbiology* **164**, 226–235 (2013).
3. Fürste, J. P. *et al.* Molecular cloning of the plasmid RP4 primase region in a multi-host-range tacP expression vector. *Gene* **48**, 119–131 (1986).
4. Lutz, R. & Bujard, H. Independent and Tight Regulation of Transcriptional Units in Escherichia Coli Via the LacR/O, the TetR/O and AraC/I1-I2 Regulatory Elements. *Nucleic Acids Res* **25**, 1203–1210 (1997).
5. Larkin, M. A. *et al.* Clustal W and Clustal X version 2.0. *Bioinformatics* **23**, 2947–2948 (2007).
6. Waterhouse, A. *et al.* SWISS-MODEL: homology modelling of protein structures and complexes. *Nucleic Acids Res* **46**, W296–W303 (2018).
7. Su, W. *et al.* Ribosome protection by antibiotic resistance ATP-binding cassette protein. *PNAS* **115**, 5157–5162 (2018).
8. Bulkley, D., Innis, C. A., Blaha, G. & Steitz, T. A. Revisiting the structures of several antibiotics bound to the bacterial ribosome. *PNAS* **107**, 17158–17163 (2010).
9. Davidovich, C. *et al.* Induced-fit tightens pleuromutilins binding to ribosomes and remote interactions enable their selectivity. *PNAS* **104**, 4291–4296 (2007).
10. Hansen, J. L. *et al.* The Structures of Four Macrolide Antibiotics Bound to the Large Ribosomal Subunit. *Molecular Cell* **10**, 117–128 (2002).
11. Ippolito, J. A. *et al.* Crystal Structure of the Oxazolidinone Antibiotic Linezolid Bound to the 50S Ribosomal Subunit. *J. Med. Chem.* **51**, 3353–3356 (2008).
12. Matzov, D. *et al.* Structural insights of lincosamides targeting the ribosome of Staphylococcus aureus. *Nucleic Acids Research* **45**, 10284–10292 (2017).
13. Svetlov, M. S. *et al.* High-resolution crystal structures of ribosome-bound chloramphenicol and erythromycin provide the ultimate basis for their competition. *RNA* **25**, 600–606 (2019).
14. Waterhouse, A. M., Procter, J. B., Martin, D. M. A., Clamp, M. & Barton, G. J. Jalview Version 2—a multiple sequence alignment editor and analysis workbench. *Bioinformatics* **25**, 1189–1191 (2009).
15. Arenz, S. *et al.* Drug Sensing by the Ribosome Induces Translational Arrest via Active Site Perturbation. *Molecular Cell* **56**, 446–452 (2014).
16. Arenz, S. *et al.* A combined cryo-EM and molecular dynamics approach reveals the mechanism of ErmBL-mediated translation arrest. *Nat Commun* **7**, (2016).
17. Beckert, B. *et al.* Structural and mechanistic basis for translation inhibition by macrolide and ketolide antibiotics. *Nat Commun* **12**, 4466 (2021).
18. Herrero del Valle, A. *et al.* Ornithine capture by a translating ribosome controls bacterial polyamine synthesis. *Nature Microbiology* **5**, 554–561 (2020).
19. van der Stel, A.-X. *et al.* Structural basis for the tryptophan sensitivity of TnaC-mediated ribosome stalling. *Nat Commun* **12**, 5340 (2021).
20. Desai, N. *et al.* Elongational stalling activates mitoribosome-associated quality control. *Science* **370**, 1105–1110 (2020).
21. Golubev, A. *et al.* Cryo-EM structure of the ribosome functional complex of the human pathogen Staphylococcus aureus at 3.2 Å resolution. *FEBS Letters* **594**, 3551–3567 (2020).
22. Li, W. *et al.* Structural basis for selective stalling of human ribosome nascent chain complexes by a drug-like molecule. *Nat Struct Mol Biol* **26**, 501–509 (2019).
